# Asymmetric ene-reduction of α,β-unsaturated compounds by F_420_-dependent oxidoreductases A (FDOR-A) enzymes from *Mycobacterium smegmatis*

**DOI:** 10.1101/2022.11.05.515281

**Authors:** Suk Woo Kang, James Antoney, David Lupton, Robert Speight, Colin Scott, Colin J. Jackson

## Abstract

The stereoselective reduction of alkenes conjugated to electron-withdrawing groups by ene-reductases has been extensively applied to the commercial preparation of fine chemicals. Although several different enzyme families are known to possess ene-reductase activity, the Old Yellow Enzyme (OYE) family has been the most thoroughly investigated. Recently, it was shown that a subset of ene-reductases belonging to the flavin/deazaflavin oxidoreductase (FDOR) superfamily exhibit enantioselectivity that is generally complementary to that seen in the OYE family. These enzymes belong to one of several FDOR subgroups that use the unusual deazaflavin cofactor F_420_. Here, we explore several enzymes of the FDOR-A subgroup, characterizing their substrate range and enantioselectivity, including the complete conversion of both isomers of citral to *(R*)-citronellel with 99% *ee*. Protein crystallography combined with computational docking has allowed the observed stereoselectivity to be mechanistically rationalized for two enzymes. These findings add further support for the FDOR and OYE families of ene-reductases being generally stereocomplementary to each other and highlight their potential value in asymmetric ene-reduction.

## Introduction

The need for pharmaceuticals and fine chemicals of high chemical and optical purity has been appreciated for the last three decades. Over this period, the environmental costs and continued viability of traditional chemical syntheses, many of which rely upon rare metals and petrochemical feedstocks that are rapidly diminishing, has raised concern.^1^ Enzymes are often seen as offering a solution to both concerns due to their high specificity and inherent renewability, coupled with the generally mild conditions in which they operate.

The reduction of alkenes is a powerful transformation in chemical synthesis, generating up to two stereogenichiral centers in a single reaction.^2, 3^ Traditional approaches for the reduction of alkenes, such as chemo/metallo-catalytic hydrogenation with transition metal complexes, have begun to be supplanted by enzymatic approaches due to their engineering plasticity and intrinsic renewability. The most well studied and widely utilized ene-reductases are those of the NAD(P)H-dependent Old Yellow Enzyme (OYE) family. These enzymes use non-covalently bound flavin mononucleotide (FMN) to transfer a hydride to the β-carbon of alkenes conjugated to electron-withdrawing groups such as ketones, aldehydes, nitriles and nitro groups.^2–4^ From the first proposal of OYEs as biocatalysts in the early 2000s,^5^ dozens of homologues have been identified and characterized, and several industrial processes now utilize OYE homologues.^6^

Ene-reductase activity has been reported in other enzyme families besides OYEs, yet these have received comparatively little attention. However, the flavin/deazaflavin oxidoreductase (FDOR) family of flavin and F_420_-dependent enzymes has received attention for their potential use as biocatalysts.^7–9^ This interest derives, in part, from their use of F_420_, a deazaflavin cofactor with an unusually low reduction potential of −340 mV, that is only sporadically distributed in nature.^10^ Known F_420_-producers include all methanogenic and some non-methanogenic archaea, many lineages of actinobacteria, certain chloroflexi, firmicutes and some proteobacteria, although it is notably absent from eukaryotes, and the key industrial microbes *Escherichia coli, Bacillus subtilis, Pseudomonas putida, Corynebacterium glutamicum*, and *Vibrio natriegens*.^11–13^ Enzymes of the FDOR superfamily are known to have physiological activity with substrates as diverse as biliverdin, menaquinone and thiopeptins.^14–16^ Several members of this family are also highly promiscuous xenobiotic reductases with activity against aflatoxins, agricultural contaminants notably recalcitrant to detoxification,^17^ as well as a variety of coumarins and triarylmethane dyes.^10, 17–19^ The activation of the nitroimidazole prodrugs pretomanid and delamanid by the deazaflavin-dependent nitroreductase (Ddn) is another key promiscuous activity of FDORs, as these represent one of the few new drug classes to treat tuberculosis.^20^

In terms of the catalytic mechanism of FDORs, the most extensively characterized enzymes are Ddn and the F_420_-dependent biliverdin reductase Rv2074. Reduced F_420_ binds to Ddn in its N1-deprotonated state that produces the more stable neutral F_420_ molecule (compared to the anionic molecule resulting from the protonated state) following hydride transfer to the substrate.^21^ The nitro group of the nitroimidazole prodrug substrate hydrogen bonds to a conserved serine residue in the active site and aligns the molecule for hydride transfer to C3 of the imidazole ring (equivalent to the β-carbon of a α,β-unsaturated nitro compound). The resulting carbocation is protonated at C2 of the imidazole ring in a net *trans*-addition of H_2_, followed by elimination of nitrous acid.^22^ in Rv2074 a conserved arginine residue is proposed to facilitate protonation of a pyrrole ring of biliverdin resulting in a resonance-stabilized carbocation at C10 to which a hydride is transferred from F_420_H_2_.^14^ Mathew *et al*. have demonstrated that the reduction of prochiral α,β-unsaturated substrates is both regio- and enantioselective.^7^ Previous studies have largely focused on characterization of the substrate scope of FDORs in terms of specific activity.^18, 19^ However, in-depth kinetic analysis and the structural basis for this selectivity remains to be determined.

In this study, we have screened a panel of the largest F_420_-dependent FDOR family, the FDOR-As, for activity with a diverse array of α,β-unsaturated compounds conjugated with a variety of electron withdrawing groups. From this initial screen we selected the two most promising enzymes, MSMEG_2027 and MSMEG_2850, for more detailed analysis. We determined the kinetic parameters and enantioselectivity with a variety of substrates. Using induced-fit computational docking, plausible Michaelis structures were obtained from which we could rationalize the observed enantioselectivity and propose a mechanism for the reduction of the substrates. Alongside recent work incorporating F_420_ biosynthesis into common microbial cell factories, such as *E. coli*,^13, 23–25^ this work should facilitate the wider adoption of FDORs as biocatalysts.

## Materials and Methods

**Chemicals** (1*R*)-(−)-myrtenal (**1a**), (+)-*cis*-myrtanol, (1*S*)-(−)-verbenone (**3a**), (*S*)-(+)-carvone (**4a**), (*R*)-(−)-carvone (**5a**), (+)-dihydrocarvone (**5b**), 2-cyclohexen-1-one (**6a**), 2-methyl-2-cyclohexen-1-one (**8a**), citral (**12a**), (*S*)-(−)-citronellal ((*S*)-**12b**), (*R*)-(+)-citronellal ((*R*)-**12b**), geraniol, nerol, 1-nitro-1-cyclohexene (**13a**), 1-cyclohexene-1-carboxaldehyde (**14a**), 1-acetyl-1-cyclohexene (**15a**), *trans*-β-nitrostyrene (**18a**), *trans*-cinnamaldehyde (**19a**), and *trans*-4-Phenyl-3-buten-2-one (**20a**) were purchased from Sigma-Aldrich (St. Louis, MO). Ketoisophorone (**2a**) was from AK Scientific (Union City, CA). 3-methyl-2-cyclohexenone (**7a**), 3-methylcyclohexanone (**7b**), 2-methylcyclohexanone (**8b**), 2-cyclopenten-1-one (**9a**), 3-methyl-2-cyclopenten-1-one (**10a**), 3-methylcyclopentanone (**10b**), 2-methyl-2-cyclopenten-1-one (**11a**), 2-methylcyclopentanone (**11b**), 1-cyclohexene-1-carboxylic acid (**16a**) and methyl-1-cyclohexene-1-carboxylate (**17a**) were provided by Enamine (Kiev, Ukraine). *trans*-myrtanol was purchased from abcr GmbH (Karlsruhe, Germany). F_420_ was obtained from a strain of *M. smegmatis* mc^2^ 4517 or *E. coli* overexpressing F_420_ biosynthesis genes as described previously.^24, 26, 27^ Other chemicals were sourced commercially at the purest grade available.

### Synthesis of *cis*-, *trans*-myrtanal (1b) and (*E*)-, (*Z*)-citral (12a)

All reactions took place under N_2_ atmosphere. For the synthesis of both *cis*-myrtanal (*cis*-**1b**) and *trans*-myrtanal (*trans*-**1b**), *cis*-myrtanol or *trans*-myrtanol (177 mg, 1.15 mmol) was dissolved in dry CH_2_Cl_2_ (60 mL), containing Dess–Martin periodinane (DMP, 600 mg, 1.41 mmol, 1.2 eq.) and stirred on ice. TLC (EtOAc/petroleum spirits = 1:3, dyed in vanillin) was used to monitor reaction progress. After 2 hours the reaction mixture was quenched with an aqueous solution of sodium thiosulfate (ca. 100–158 g/L, Na_2_S_2_O_3_), and the organic phase was washed with NaHCO_3_ (ca. 100 g/L-saturated), water, brine, and then dried over Na_2_SO_4_. The filtered mixture was concentrated under reduced pressure. Using flash chromatography (SiO_2_, diethyl ether/n-hexane = 4:96), the residue was purified to yield *cis*-myrtanal (*cis*-**1b**, 160.3 mg, 90.6 %) and *trans-* myrtanal (*trans*-**1b**, 155.7 mg, 88.0 %) as colorless oil. Spectral data matched literature values.^28^

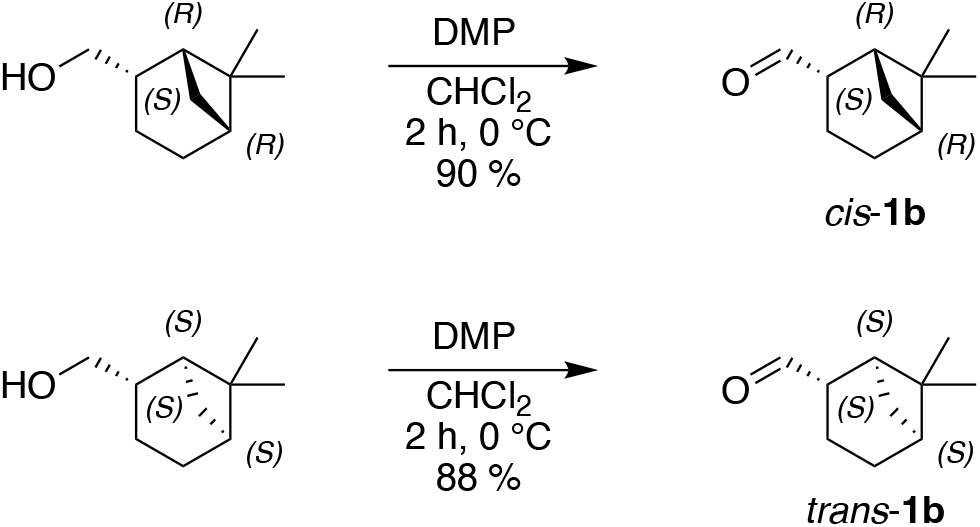

For the synthesis of *trans*-**12a**, *cis*-**12a**, either geraniol or nerol (190 mg, 1.23 mmol) was dissolved in dry CH_2_Cl_2_ (60 mL) containing DMP (650 mg, 1.53 mmol, 1.25 eq.) for on ice for 2 hours. Following the same workup as above afforded geranial (*trans*-**12a**, 170.2 mg, 89.6 %) and neral (*cis*-**12a**, 164.5 mg, 86.6 %) as colorless oils. The products are confirmed using GC-MS and/or NMR with comparison to literature values.^7^

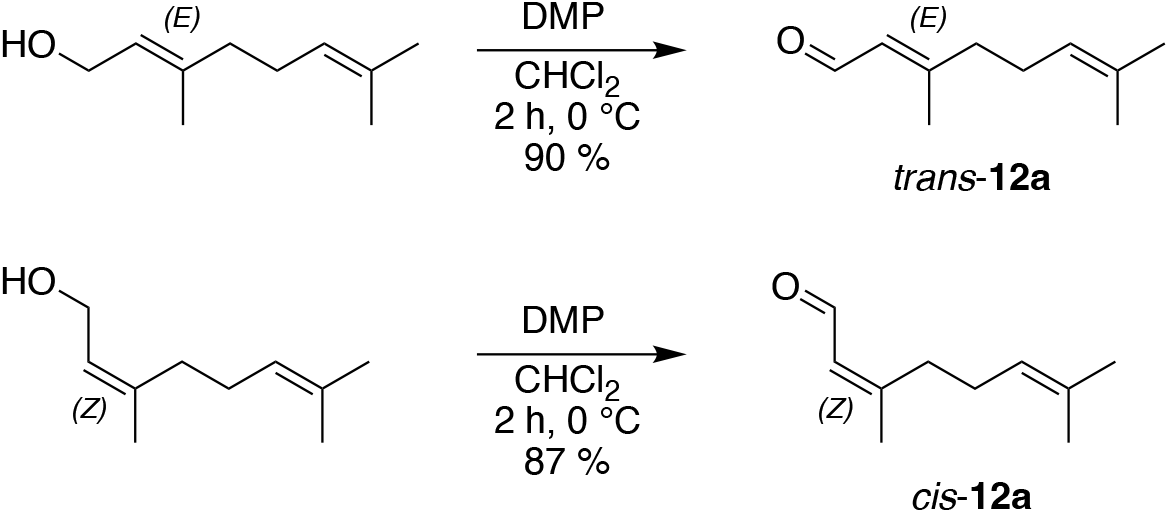

### Cloning, expression, and purification of proteins

Plasmids expressing F_420_H_2_-dependent FDORs and F_420_-dependent glucose-6-phosphate dehydrogenase (Fgd) have been described previously^10, 17, 29, 30^ and are detailed in **Table S1**. For crystallographic studies, the codon optimized MSMEG_2850 gene with a TEV cleavage site was synthesized (IDT) and cloned into pETMCSIII by Gibson assembly as described previously.^10^ Transformants were plated on LB agar plates containing 100 µg mL^−1^ ampicillin and incubated at 37 °C overnight. Starter cultures were inoculated from single colonies and grown in 10 mL of LB media supplemented with ampicillin at 37 °C overnight. The starter cultures were transferred to 1 L of modified auto-induction TB 2.0 media, containing 5 g L^−1^ yeast extract, 20 g L^−1^ tryptone, 85.5 mM NaCl, 22 mM KH_2_PO_4_, 42 mM Na_2_HPO_4_, 0.6% glycerol, 0.05% glucose, 0.2% lactose and 100 µg mL^−1^ ampicillin. The large-scale cultures were grown at 30 °C with shaking at 180 rpm overnight. Cells were pelleted by centrifugation at 5 000*g* for 15 min at 4 °C, resuspended in 40 mL of lysis buffer (50 mM NaH_2_PO_4_, 300 mM NaCl, 25 mM imidazole, pH 8.0) adding 1 µL of turbonuclease (Sigma) and lysed by sonication using an Omni Sonicator Ruptor 400 (2 × 3 min at 60% power). The soluble fraction was separated by further centrifugation at 13 000 x *g* for 1 h at 4 °C and the supernatant was loaded onto a 5 mL HisTrap FF NiNTA column (GE Healthcare) equilibrated with lysis buffer. The column was washed with 5 column volumes of lysis buffer containing 25 mM imidazole, followed by 5 column volumes of buffer containing 40 mM imidazole. Purified protein was eluted with buffer containing 250 mM imidazole. Purified proteins were dialysed overnight against 4 L of assay buffer (50 mM Tris, 300 mM NaCl, pH 8) supplemented with 10% glycerol at 4 °C. Aliquots of purified protein were flash frozen and stored at −80 °C until use.

For crystallography, the His_6_-tag was cleaved with TEV protease (expressed and purified in-house^31^) in 50 mM Tris, 150 mM NaCl, 10 mM DTT, pH 8.0 overnight at room temperature. Following a subtractive NiNTA pass to remove the TEV protease, the samples were concentrated and purified on a HiLoad 16/60 Superdex 75 pg (GE Healthcare) equilibrated with 20 mM HEPES, 150 mM NaCl, pH 7.5. To generate holo-complexes with F_420_, samples were incubated with an excess of F_420_ at room temperature for at least 1 hour prior to size-exclusion.

### Protein crystallography

Following size exclusion MSMEG_2850 was exchanged into 20 mM HEPES, 50 mM NaCl pH 7.5 and concentrated to 3.6 mM (∼50 mg mL^−1^). Initial screens were conducted using SG1 (Molecular Dimensions), PEG/ION, and HT sparse matrix screens (Hampton Research). Drops containing1 μL purified protein and 1 μL of mother liquor were equilibrated against 100 µL mother liquor at 18 °C. Optimised conditions for the crystals used for data collection were 1.3 M potassium phosphate dibasic, 0.7 M sodium phosphate monobasic, pH 6.9. Crystals were soaked in cryobuffer (1.3 M potassium phosphate dibasic, 0.7 M sodium phosphate monobasic, 30% glycerol, pH 6.9) for X-ray diffraction.

### Data acquisition and structure refinement

X-ray diffraction data were collected at the Australian Synchrotron at the MX2 beamline. Diffraction data were integrated using XDS^32^ and scaled using AIMLESS^33^ in the CCP4 suite.^34^ Structures were solved using molecular replacement with PHASER^35^ with 4Y9I as the search model. Model building was performed in COOT.^36^ Refinements were performed using REFMAC^37^ and phenix.refine.^38^

### Enzyme activity assays

F_420_ was reduced enzymatically with 0.2–1.0 μM Fgd and excess G6P in assay buffer that had been degassed under nitrogen. Fgd was removed by diafiltration (Amicon Ultra, 10 kDa, Merck Millipore) and F_420_H_2_ was used immediately after preparation.

For the initial broad activity screen of the FDORs, F_420_ was reduced overnight in 50 mM Tris, 150 mM NaCl, pH 7.5 and separated from Fgd by diafiltration. 1 µM FDOR was preincubated with 10 µM F_420_H_2_ for 5 minutes. Assays were initiated by addition of 100 µM substrate. Activity was determined fluorometrically by following the reoxidation of F_420_H_2_ (λ_ex_ = 420 nm, λ_em_ = 470 nm) on a SpectraMAX microplate spectrofluorimeter (Molecular Devices)^29^. Assays were conducted at room temperature and each combination of enzyme and substrate was tested in duplicate. To explore the relationship between sequence, chemical structure, and activity, protein sequences were aligned with the T-Coffee web server using the Expresso method^39^ and a nearest-neighbour tree constructed with MEGA X.^40^ For the substrates, MACCS fingerprints were generated using Canvas (Schrodinger LLC, NY) and clustered hierarchically using Tanimoto similarity.

For more detailed kinetic studies, assays were performed in 50 mM Tris, 300 mM NaCl, pH 8.0. Specific activities were measured in 100 μL reactions containing 0.1 µM FDOR, 20 µM F_420_H_2_ and 250 µM substrate. 100 mM stock solutions of substrates were prepared in methanol, except ketoisophorone (**2a**) which was dissolved in the assay buffer. Kinetic data were obtained as above with substrates varied from 3.08 µM to 20 mM. Apparent steady-state kinetic parameters were calculated by fitting to the standard Michealis–Menten equation (1) or Michaelis–Menten with substrate inhibition equation (2) as appropriate using GraphPad Prism v.8.3 (GraphPad Software Inc., La Jolla, CA). Assays were conducted in duplicate.

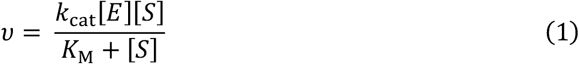

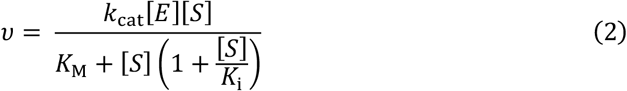

### GC/MS analysis

Enzyme reactions were performed in Eppendorf tubes containing 72.5 µM F_420_, 2.9 µM FDOR, 1.6 mM substrate, 6.5 mM G6P and 1.4 µM Fgd in a final volume of 155 μL. Reactions were extracted with an equal volume of ethyl acetate containing 1 mM mesitylene as an internal standard. Samples were analysed using GC-MS (7890A GC system, 5975C MSD, Agilent, Santa Clara, CA). To measure conversions, samples were separated on an Agilent HP-1 column (30 m × 0.25 mm × 0.25 μm) with the following settings: Injection temp: 280 °C; Oven program: 50 °C for 2 min; 4 °C min^−1^ until 100 °C for 0 min; 10 °C min^−1^ until 250 °C for 10 mins. Nitrogen was used as a carrier gas (1.2 mL min^−1^). 1 μL of sample was injected into the GC and the split ratio was 50:1. Substrate and products were identified using the NIST MS spectrum library and the identification was confirmed with commercial or synthesized standards. Retention times are listed in **Table S2**. Chromatograms are shown in **Figure S1**.

Enzymatic reductions of prochiral substrates (**1a, 2a, 4a, 5a, 7a, 8a, 10a**–**12a**) were performed as above. Time course analyses were conducted by sampling 155 μL from reaction mixture at 0.5, 1.0, 2.0 and 24 h after initiation of the reaction (**Figure S2**). Each sample was extracted with an equal volume of ethyl acetate containing 1 mM mesitylene and analyzed by chiral GC-FID using an Agilent 7890A system (**Figure S1**). The absolute configurations of reaction products were determined by comparing the retention time with standard compounds of known configuration or with previously reported retention indices^7, 41–43^. Agilent CycloSil B column (30 m × 0.25 mm × 0.25 μm) was used for the separation of substrates **1a, 4a, 5a, 7a, 10a, 11a** and **12a** and Agilent CP-Chirasil-Dex CB column (25 m × 0.32 mm × 0.25 µm) for substrates **2a** and **8a**. Injection temperature of 250 °C and a 50:1 split ratio employed. Separation conditions and retention times are described in **Table S3**.

### Computational docking

To dock substrates into the active sites of MSMEG_2027 and MSMEG_2850, the structures were superimposed in PyMOL (Schrodinger LLC, NY) and the F_420_-4 molecule of the holo MSMEG_2027 structure (PDB: 6WTA) was copied into the MSMEG_2850 structure. The rotamer of R48 was adjusted to remove a clash with F_420_. The loop between α1 and β1 in the structure of MSMEG_2850 was rebuilt using Prime loop refinement (Schrodinger LLC, NY) as it showed substantial deviation in Cα rmsd compared to structures of homologous enzymes due to its involvement in a domain-swapped dimer interface. The models of both complexes are shown in **Figure S3**. LigPlot plus was used to analyze interactions with F_420_ and active site residues (**Figure S3**). Both structures were parameterized with the OPLS3e forcefield and subjected to energy minimization using Desmond (Schrodinger LLC, NY) with the default settings.^44^ Following minimization, waters were removed from the structures, and substrates parameterized with OPLS3e were docked into the active sites using the induced fit protocol in Maestro 2019-01 release (Schrodinger LLC, NY).

### Circular dichroism thermostability assays

Circular dichroism experiments were performed using a ChiraScan spectrometer (Applied Photophysics) with a Quantum temperature controller (NorthWest). FDORs and F_420_ were prepared in 10 mM phosphate buffer (pH 7.5) and then mixed and diluted to the final samples containing 11.25 µM of FDOR and various concentrations of F_420_ (0–112.5 µM). All samples were incubated in a 400 µL quartz cuvette (path length = 1 mm) for 10 min at 20 °C before analysis. Far-UV (190–260 nm) spectra were recorded with a 1 nm slit width over the temperature range from 20 °C to at least 90 °C in 2 °C increments (1 °C min^−1^), with measurements at each temperature following a 1 min equilibration. Measurements were conducted in duplicate. Thermostability of FDORs was monitored by single wavelength denaturation curve at 218 nm and the melting temperature (*T*_M_) was calculated from the midpoint of the denaturation curve using Boltzmann sigmoidal in GraphPad Prism v8.3 (GraphPad Software Inc., La Jolla, CA).

## Results

### Substrate specificity of the FDOR-As

We first heterologously expressed and purified seven FDOR-A enzymes from *M. smegmatis* (MSMEG_2850, MSMEG_2027, MSMEG_5998, MSMEG_5215, MSMEG_3909), *Mycobacterium tuberculosis* (Rv3547), and *Rhodococcus erythropolis* (RER_09240) in *E. coli*. These purified enzymes were then screened for activity against a broad panel of α,β-unsaturated compounds in a high throughput semi-quantitative screen (**Figure 1**). We found that all enzymes tested had detectable activity with multiple substrates, consistent with previous showing broad substrate specificity in the FDOR-As.^18, 19, 30^ The only homologue within this set from a non-mycobacterial source (RER_09240) displayed broad but low-level activity, comparable to that of MSMEG_5998 (**Figure 1**). Rv3547 (Ddn) showed moderately high activity with a broad range of substrates but does not express in soluble form in *E. coli* without a maltose-binding protein solubility tag.^45, 46^ From our initial screen, we selected MSMEG_2027 and MSMEG_2850 for further characterization based on their broad substrate coverage and high specific activities, in addition to their high soluble expression in *E. coli* without the need for a solubility tag.

**Figure 1.**
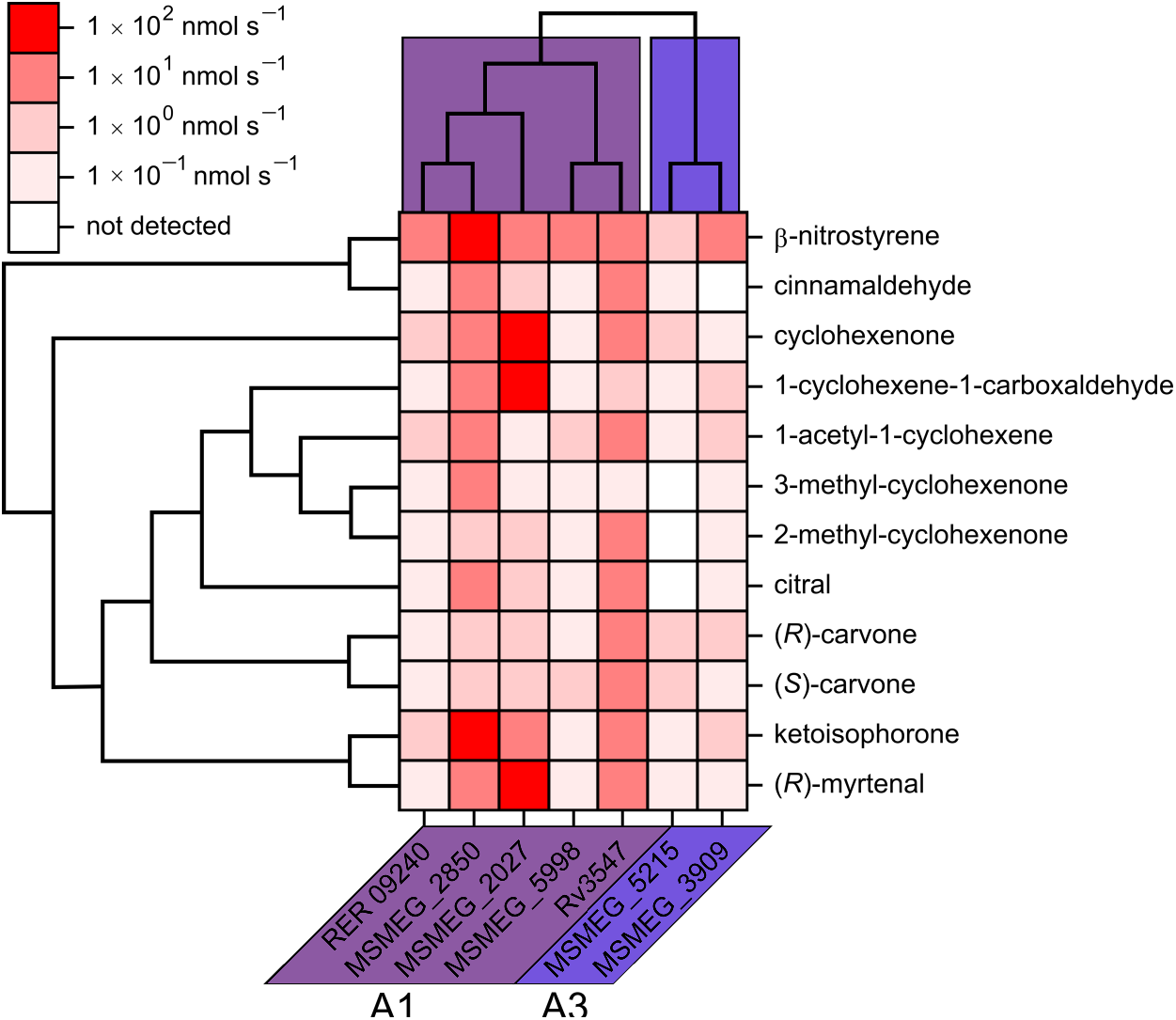
Results of initial screen of FDORs with a panel of α,β-unsaturated compounds. Assays were conducted with 1 µM enzyme, 10 µM F_420_H_2_, and 100 µM substrate. Sequences were aligned using the T-Coffee web server with Expresso and a nearest-neighbor tree generated with MEGA X. Fingerprints were generated using MACCS keys and hierarchically clustered based on Tanimoto similarity using Canvas. Activity expressed as nmoles of F_420_H_2_ oxidized per second.

### Organic cosolvent tolerance

Many substrates suffer from low aqueous solubility and organic cosolvents are often employed to overcome this. We therefore measured activity in the presence of different organic cosolvents at 10% (v/v) (**Figure 2**) using **1a** and **2a** as substrates for MSMEG_2027 and MSMEG_2850, respectively, based on the high activity identified in the initial screen. Both enzymes followed the same trend in cosolvent tolerance, with protic solvents being better tolerated than aprotic ones. Acetone and acetonitrile were found to be particularly inhibitory, both resulting in approximately 50% and 20% residual activity for MSMEG_2027 and MSMEG_2850, respectively. DMSO was well tolerated by MSMEG_2027 and was the least inhibitory aprotic solvent tested. Methanol was the least inhibitory overall for both enzymes with 89% and 67% residual activity for MSMEG_2027 and MSMEG_2850, respectively. The effect of varying methanol concentration from 0–25% was investigated (**Figure 2**). MSMEG_2027 consistently retained a greater proportion of activity than MSMEG_2850 at a given methanol concentration. Both enzymes retained greater than 90% activity in the presence of 2.5% methanol, with greater than 50% activity retained at 15% (v/v) for MSMEG_2850 and 25% (v/v) for MSMEG_2027. Based on these results methanol was selected as the cosolvent for all subsequent activity assays.

**Figure 2:**
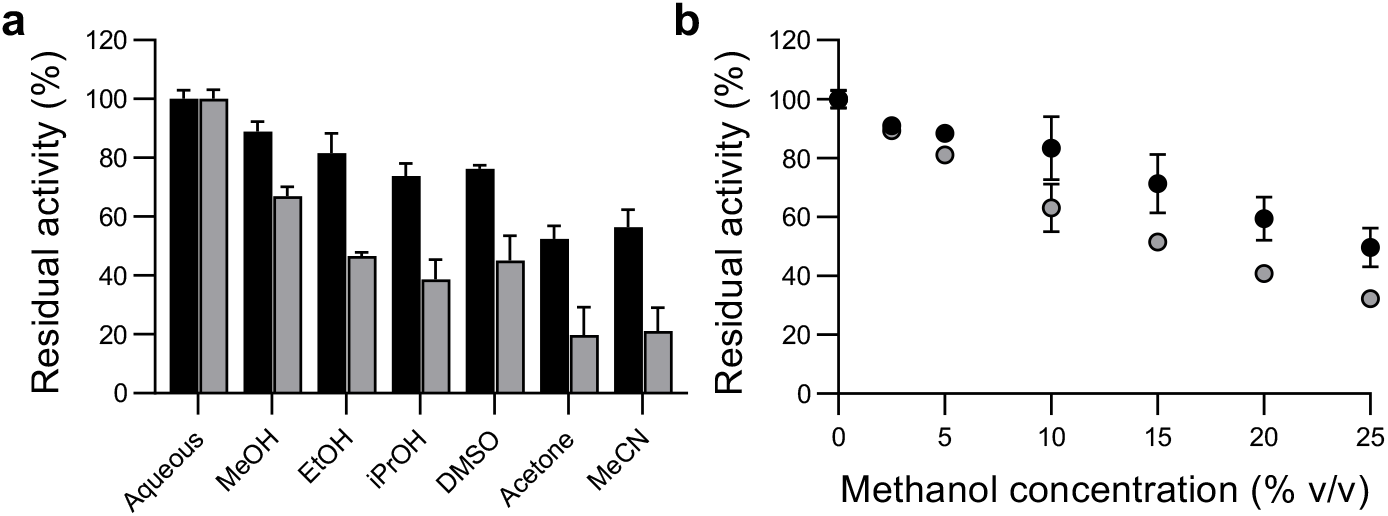
Organic cosolvent tolerance of MSMEG_2027 and MSMEG_285. (a) Residual activity in the presence of 10% v/v of the indicated organic solvent in 50 mM Tris-HCl pH 8.0 with 20 µM F_420_, 25 µM substrate (**1a** and **2a** for MSMEG_2027 and MSMEG_2850, respectively) and 0.1 µM enzyme. (b) Residual activity with increasing concentrations of methanol from 0–25%. Black bars and circles, MSMEG_2027; grey bars and circles, MSMEG_2850.

With assay conditions optimized, we measured the specific activity of MSMEG_2027 and MSMEG_2850 with 20 substrates containing aldehyde, ketone, carboxylate, ester, and nitro groups (**Table 1**). Substrate conversion was confirmed by GC-MS (**Table S2, Figure S1**). Consistent with the results of the initial screen, both enzymes accepted a range of substrates that includes several typical OYE substrates (**Table 1**). We also obtained steady-state kinetic parameters for each substrate and enzyme pair (**Table 2; Figure S2**). Specific activities varied from 0.5 to 84.4 μmol min^−1^ μmol^−1^ enzyme for MSMEG_2027 with **17a** and **6a**, respectively and 0.3 to 39.9 μmol min^−1^ μmol^−1^ enzyme for MSMEG_2850 with **8a** and **10a**, and **18a**, respectively. Notably, MSMEG_2027 displayed high activity with (*R*)-myrtenal **1a** which has hitherto received little study as a substrate for ene-reductases.^47^

**Table 1:**
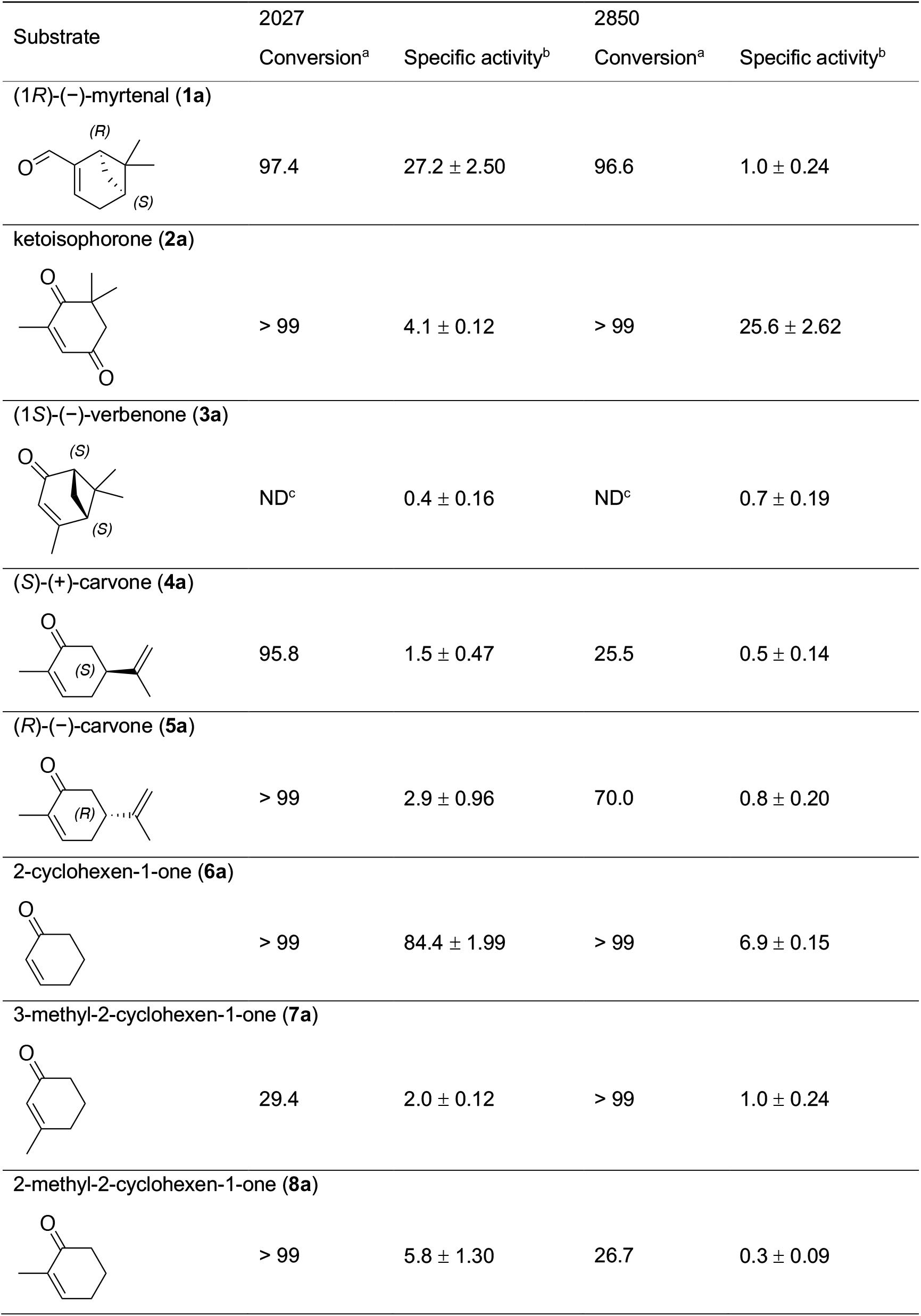
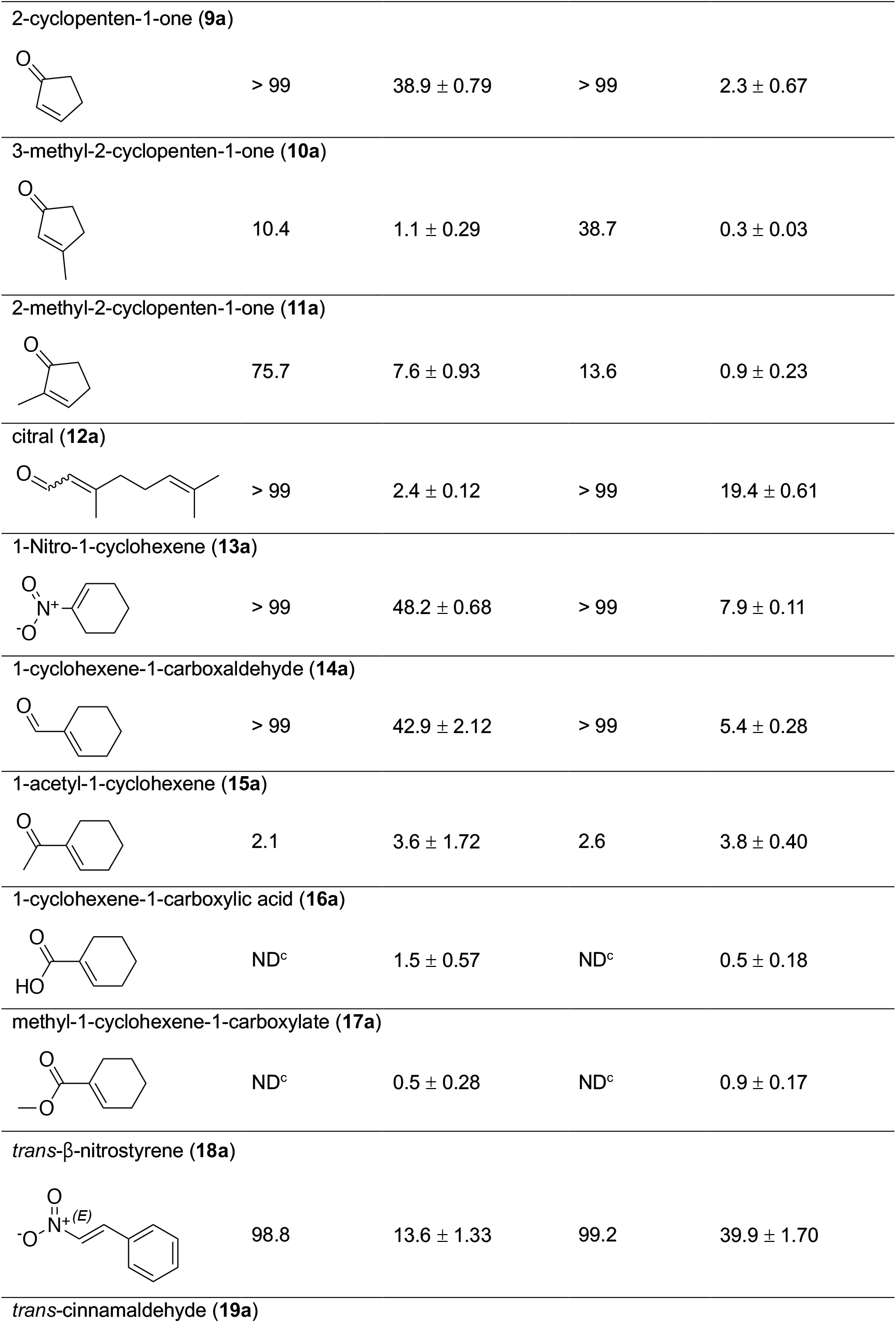
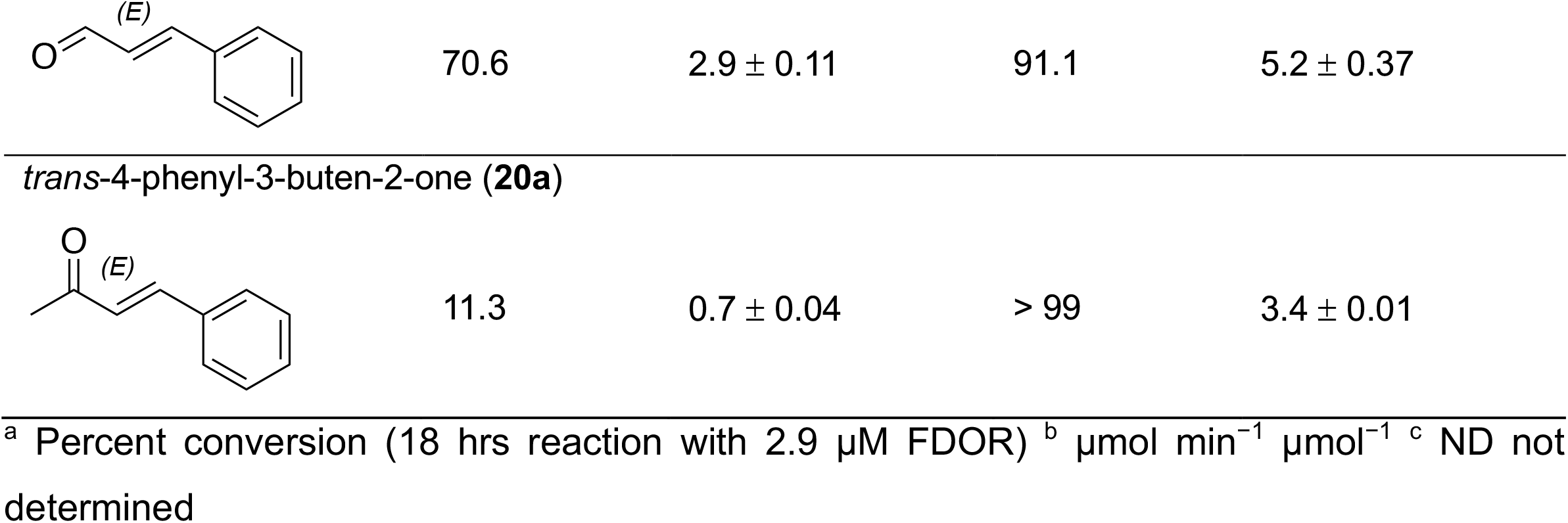
Conversions and specific activities for substrates **1a**–**20a**

| Substrate | 2027 |  | 2850 |  |
| --- | --- | --- | --- | --- |
|  | Conversion<sup>a</sup> | Specific activity<sup>b</sup> | Conversion<sup>a</sup> | Specific activity<sup>b</sup> |
| (1*R*)-(-)-myrtenal (**1a**) |  |  |  |  |
|  | 97.4 | 27.2 ± 2.50 | 96.6 | 1.0 ± 0.24 |
| ketoisophorone (**2a**) |  |  |  |  |
|  | > 99 | 4.1 ± 0.12 | > 99 | 25.6 ± 2.62 |
| (1*S*)-(-)-verbenone (**3a**) |  |  |  |  |
|  | ND<sup>c</sup> | 0.4 ± 0.16 | ND<sup>c</sup> | 0.7 ± 0.19 |
| (*S*)-(+)-carvone (**4a**) |  |  |  |  |
|  | 95.8 | 1.5 ± 0.47 | 25.5 | 0.5 ± 0.14 |
| (*R*)-(-)-carvone (**5a**) |  |  |  |  |
|  | > 99 | 2.9 ± 0.96 | 70.0 | 0.8 ± 0.20 |
| 2-cyclohexen-1-one (**6a**) |  |  |  |  |
|  | > 99 | 84.4 ± 1.99 | > 99 | 6.9 ± 0.15 |
| 3-methyl-2-cyclohexen-1-one (**7a**) |  |  |  |  |
|  | 29.4 | 2.0 ± 0.12 | > 99 | 1.0 ± 0.24 |
| 2-methyl-2-cyclohexen-1-one (**8a**) |  |  |  |  |
|  | > 99 | 5.8 ± 1.30 | 26.7 | 0.3 ± 0.09 |
| 2-cyclopenten-1-one (**9a**) |  |  |  |  |
| | > 99 | $38.9 \pm 0.79$ | > 99 | $2.3 \pm 0.67$ |
| 3-methyl-2-cyclopenten-1-one (**10a**) |  |  |  |  |
| | 10.4 | $1.1 \pm 0.29$ | 38.7 | $0.3 \pm 0.03$ |
| 2-methyl-2-cyclopenten-1-one (**11a**) |  |  |  |  |
| | 75.7 | $7.6 \pm 0.93$ | 13.6 | $0.9 \pm 0.23$ |
| citral (**12a**) |  |  |  |  |
| | > 99 | $2.4 \pm 0.12$ | > 99 | $19.4 \pm 0.61$ |
| 1-Nitro-1-cyclohexene (**13a**) |  |  |  |  |
| | > 99 | $48.2 \pm 0.68$ | > 99 | $7.9 \pm 0.11$ |
| 1-cyclohexene-1-carboxaldehyde (**14a**) |  |  |  |  |
| | > 99 | $42.9 \pm 2.12$ | > 99 | $5.4 \pm 0.28$ |
| 1-acetyl-1-cyclohexene (**15a**) |  |  |  |  |
| | 2.1 | $3.6 \pm 1.72$ | 2.6 | $3.8 \pm 0.40$ |
| 1-cyclohexene-1-carboxylic acid (**16a**) |  |  |  |  |
| | ND<sup>c</sup> | $1.5 \pm 0.57$ | ND<sup>c</sup> | $0.5 \pm 0.18$ |
| methyl-1-cyclohexene-1-carboxylate (**17a**) |  |  |  |  |
| | ND<sup>c</sup> | $0.5 \pm 0.28$ | ND<sup>c</sup> | $0.9 \pm 0.17$ |
| *trans*-β-nitrostyrene (**18a**) |  |  |  |  |
| | 98.8 | $13.6 \pm 1.33$ | 99.2 | $39.9 \pm 1.70$ |
| *trans*-cinnamaldehyde (**19a**) |  |  |  |  |
| | 70.6 | $2.9 \pm 0.11$ | 91.1 | $5.2 \pm 0.37$ |
| *trans*-4-phenyl-3-buten-2-one (**20a**) |  |  |  |  |
| | 11.3 | $0.7 \pm 0.04$ | > 99 | $3.4 \pm 0.01$ |
| <sup>a</sup> Percent conversion (18 hrs reaction with 2.9 μM FDOR) <sup>b</sup> μmol min<sup>-1</sup> μmol<sup>-1</sup> <sup>c</sup> ND not determined |  |  |  |  |

**Table 2:**
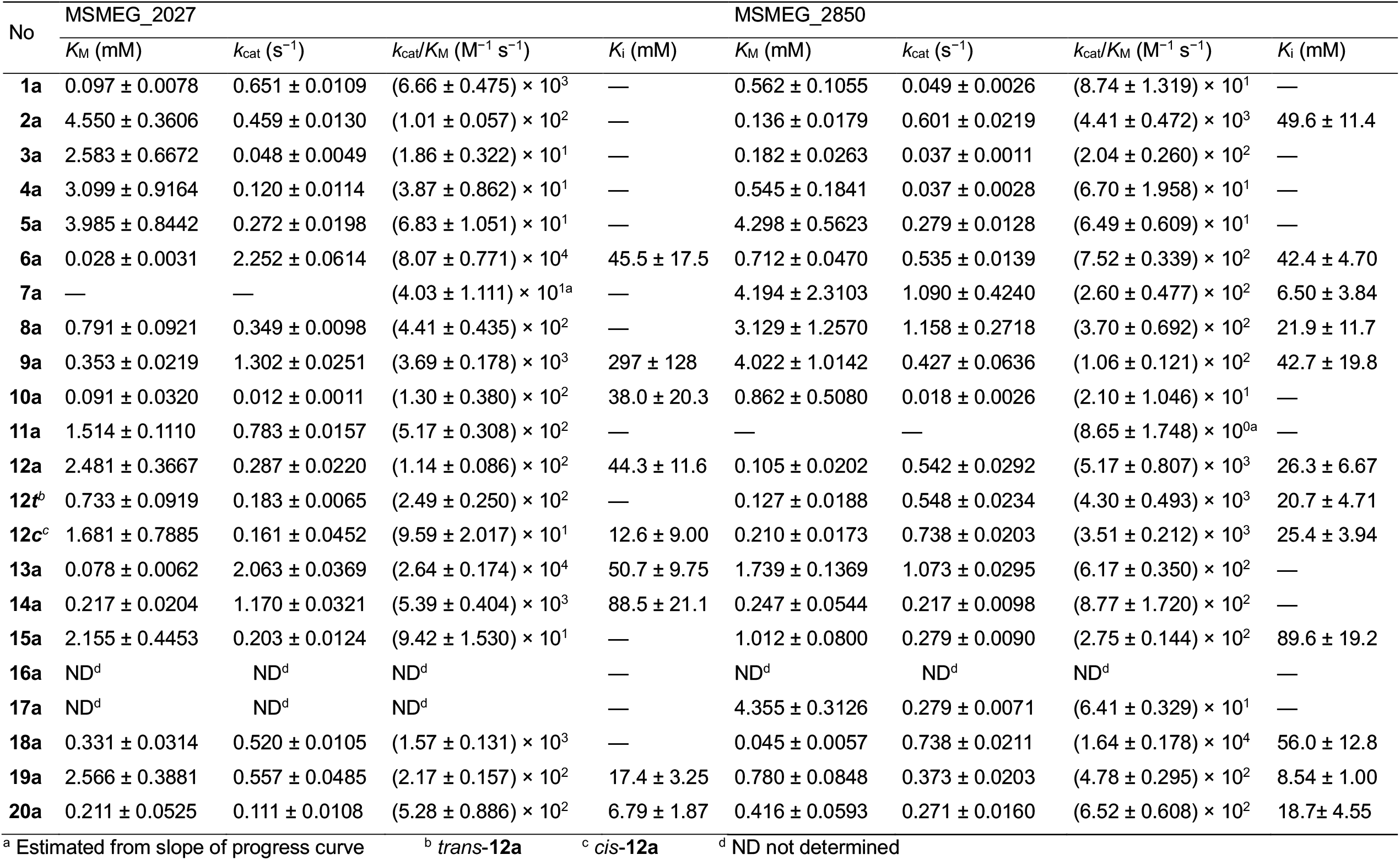
Kinetic parameters for substrates **1a**–**20a**

MSMEG_2850 had high activity with **2a**, while both enzymes showed a preference for **5a** over **4a** as substrates (**Table 1**). Interestingly, both enzymes displayed higher affinity but lower turnover of **4a** compared to **5a** under steady-state conditions (**Table 2)**. Both enzymes preferred unsubstituted endocyclic enones **6a** and **9a**, to the 2-methyl and 3-methyl derivatives. Addition of a methyl group at the β-position (**7a**) to the 2-cyclohexenone scaffold was less well tolerated by MSMEG_2027 than addition to the α-position (**8a**). Interestingly, the reverse was observed for MSMEG_2850. Based on specific activity and catalytic efficiency (**Tables 1**,**2**), methyl substitution at the β-position (**10a**) of 2-cyclopentenone was more inhibitory for both enzymes than that at α-position (**11a**) and goes against the general trend seen with OYE homologues.^43, 48, 49^ However, a higher conversion rate of **11a** compared to **10a** was observed for MSMEG_2850. The fluorescence detection assay did not always quantitatively agree with the conversion as measured by GC, especially when specific activity was very low (e.g., **3a, 16a**, and **17a**). This was probably due to the signal:noise ratio at low levels of activity.

The specific activity of **6a** and **9a** was approximately 10-fold higher with MSMEG_2027 compared to MSMEG_2850, while the specific activity of **6a** is higher than that of **9a** with both enzymes (2.2- and 3-fold for MSMEG_2027 and MSMEG_2850, respectively, (**Table 1**). Steady-state kinetics provided insight into the large difference in activity with **6a**. MSMEG_2850 had approximately 4-fold lower *k*_cat_, 25-fold higher *K*_M_ (**Table 2**). In contrast, the specific activity of citral **12a** was 8-fold higher with MSMEG_2850 than with MSMEG_2027 (**Table 1**). GC analysis showed that the reaction was regioselective as only the activated double bond at C2 was reduced while the non-activated double bond at C6 was untouched. The activity of the exocyclic cyclohexene series correlated with the strength of the EWG; high (48.2–5.4 μmol min^−1^ μmol^−1^ enzyme) for nitro group (**13a**) and aldehyde (**14a**), moderate (3.6– 3.8 μmol min^−1^ μmol^−1^ enzyme) for ketone (**15a**) and low (0.5–1.5 μmol min^−1^ μmol^−1^ enzyme) for carboxylate (**16a**) and ester (**17a**) derivatives, consistent with trends seen in the OYE family.^50, 51^ This trend was also observed with a series of cinnamic acid derivatives. Cinnamaldehyde (**19a**) exhibited 4.6- and 7.6-fold lower activity with MSMEG_2027 and MSMEG_2850, respectively, compared to nitrostyrene (**18a**), while 4-Phenyl-3-buten-2-one (**20a**) was 4.1- and 1.5-fold less active than cinnamaldehyde (**19a**). Cinnamic acid and its methyl ester were not turned over by either enzyme (data not shown).

Steady-state kinetic parameters allowed further insight into the differences in activity between substrates. Comparing **1a** with **14a**, that both contain an exocyclic aldehyde, we found that the *K*_M_ for **1a** is lower than that of **14a** with MSMEG_2027 while the converse is true with MSMEG_2850 (**Table 2**). The 2.3-fold lower *K*_M_ of **1a** is largely compensated for by a 1.8-fold lower *k*_cat_ compared to **14a** with MSMEG_2027, yielding comparable catalytic efficiencies, with *k*_cat_/*K*_M_ = (6.66 ± 0.475) and (5.39 ± 0.404) × 10^3^ M^−1^ s^−1^, respectively (**Table 2**).

Comparing **3a, 7a** and **10a**, each bearing a methyl group on the β-carbon, it is interesting that the *K*_M_ of the smaller **10a** (0.091 ± 0.032 and 0.86 ± 0.51 mM, respectively for MSMEG_2027 and MSMEG_2850) is smaller than that of the larger **7a** (which we could not measure for MSMEG_2027, 4.19 ± 2.31 mM for MSMEG_2850, table 2) Interestingly, saturation of **7a** does not appear to occur even at 20 mM for MSMEG_2027, while for MSMEG_2850 *k*_cat_ is relatively high (1.090 ± 0.424 s^−1^) however the *K*_i_ is comparable to the *K*_M_ (6.50 ± 3.84 and 4.194 ± 2.310 mM, respectively) indicating severe substrate inhibition. This accounts for the difference in conversion of **7a** between MSMEG_2027 and MSMEG_2850 (**Table 1**). Conversely MSMEG_2027 showed substrate inhibition with **10a** while no substrate inhibition was observed for **3a** with either enzyme. It could be that the additional steric bulk and/or rigidity imposed on **3a** compared to **7a** by its substituents prevent it from binding in a competitive manner. These substrates typically had by low values for *k*_cat_ compared to substrates bearing methyl groups on the α-carbon.

Compounds **2a, 4a, 5a, 8a** and **11a** all bear a methyl group on the α-carbon and generally exhibited modest activities, except for **2a** with MSMEG_2850. Of the two enantiomers of carvone both enzymes showed a preference for **5a** over **4a**. Interestingly MSMEG_2027 had comparable *K*_M_ values for **4a** and **5a**, while for MSMEG_2850 the *K*_M_ for **4a** was nearly 8-fold lower than that of **5a** (table 2). Concomitantly the *k*_cat_ of **4a** was 7.5-fold lower than that of **5a**, leading to comparable catalytic efficiencies [(6.70 ± 1.958) and (6.49 ± 0.609) × 10^1^ M^−1^ s^−1^, respectively, **Table 2**]. Similarly, the catalytic efficiencies of **8a** and **11a** with MSMEG_2027 are comparable while MSMEG_2850 used **8a** more effectively than **11a**.

Citral **12a** and its individual isomers neral and geranial (**12c** and **12t** in **Table 2**) also show different activity profiles. MSMEG_2850 has comparable catalytic efficiency for both isomers as with the (3:2) mixture found in citral (*k*_cat_/*K*_M_ between 3.51 × 10^3^ and 5.17 × 10^3^ M^−1^ s^−1^, **Table 2**) while MSMEG_2027 had a 2.3-fold lower catalytic efficiency for **12c** compared to **12t**, largely due to a 2.3-fold higher *K*_M_ value for the later compared to the former (**Table 2**).

Patterns of substrate inhibition differed substantially between the two enzymes, with **10a, 13a** and **14a** inhibiting MSMEG_2027 but not MSMEG_2850, **2a, 7a, 8a, 12t, 15a**, and **18a** inhibiting MSMEG_2850 but not MSMEG_2027, and **6a, 9a, 12a, 12c, 19a** and **20a** inhibiting both enzymes. **1a, 3a**–**5a**, and **11a** showed no detectable substrate inhibition with either enzyme. Inhibition was particularly strong with MSMEG_2850 and substrates **7a** and **19a** in which the *K*_i_ was found to be 1.5- and 11-fold the *K*_M_, respectively (**Table 2**), as well as MSMEG_2027 with **12c** (7.5-fold higher *K*_i_ than *K*_M_).

### Stereochemical analysis

To determine if reduction of prochiral substrates occurred stereoselectively we analyzed the products of bioreductions by chiral GC (**Table 3**). MSMEG_2027 displayed excellent conversion (>99%) in 30 min with moderate diastereoselectivity (82.3% *de*) for (*R*)-myrtenal **1a**. MSMEG_2850 required an extended reaction time (24 h) to convert 98% of **1a** with only 68.6% *de*. In contrast, ketoisophorone **2a** was completely converted by MSMEG_2850 within 30 min to yield (*S*)-levodione **2b** (99% *ee*), while 2 hours was required for complete conversion by MSMEG_2027 with a lower optical purity (90.1% *ee*), in part the lower selectivity was due to racemization of the product as observed in the time-course chromatogram (**Figure 3**).^41^ The conversion rates are consistent with the results of specific activities and similarly applied to the other substrates (**Table 1**).

**Table 3:**
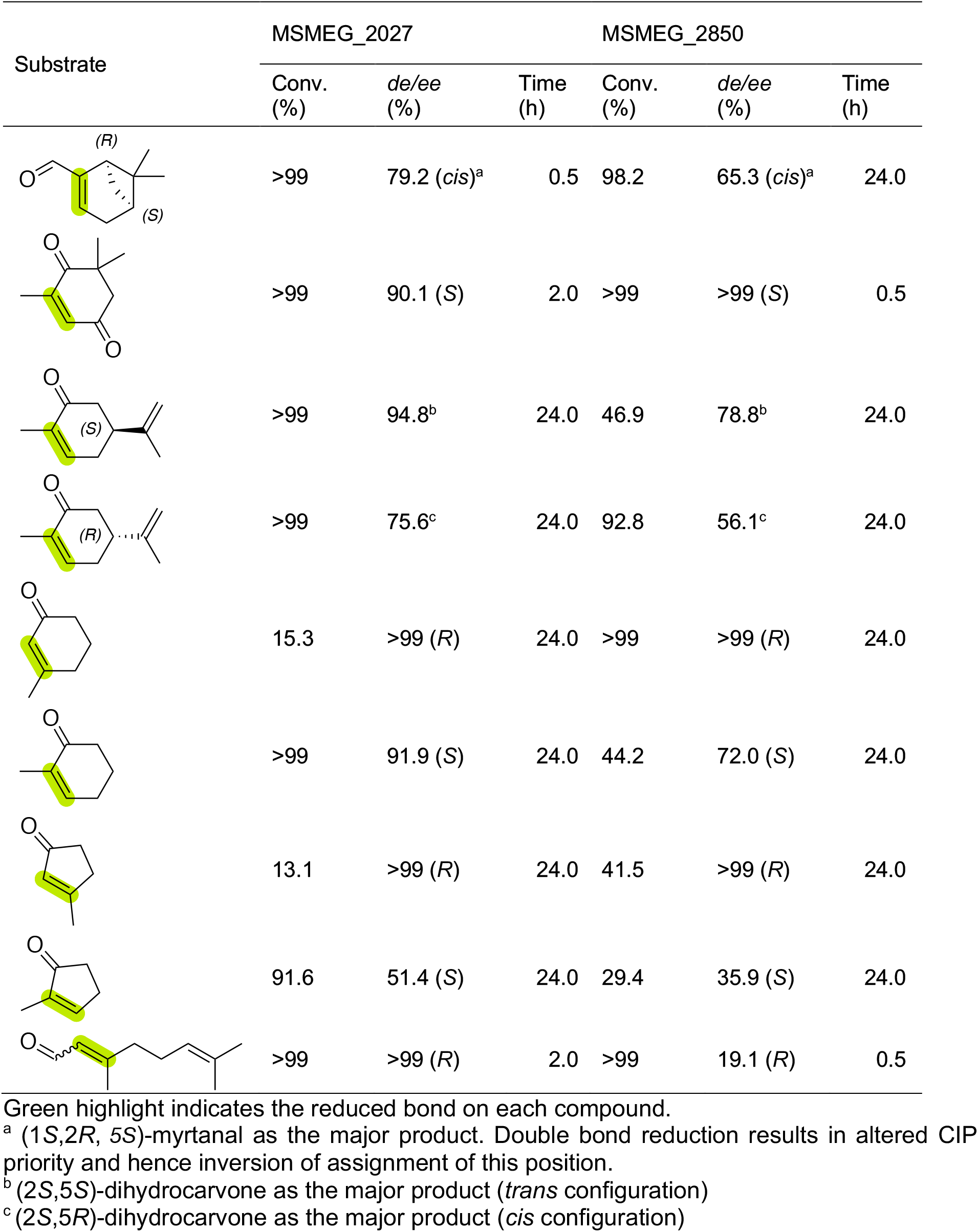
Stereochemical outcome of FDOR biotransformation

| Substrate | MSMEG_2027 |  |  | MSMEG_2850 |  |  |
| --- | --- | --- | --- | --- | --- | --- |
|  | Conv. (%) | de/ee (%) | Time (h) | Conv. (%) | de/ee (%) | Time (h) |
|  | >99 | 79.2 (*cis*)<sup>a</sup> | 0.5 | 98.2 | 65.3 (*cis*)<sup>a</sup> | 24.0 |
|  | >99 | 90.1 (S) | 2.0 | >99 | >99 (S) | 0.5 |
|  | >99 | 94.8<sup>b</sup> | 24.0 | 46.9 | 78.8<sup>b</sup> | 24.0 |
|  | >99 | 75.6<sup>c</sup> | 24.0 | 92.8 | 56.1<sup>c</sup> | 24.0 |
|  | 15.3 | >99 (R) | 24.0 | >99 | >99 (R) | 24.0 |
|  | >99 | 91.9 (S) | 24.0 | 44.2 | 72.0 (S) | 24.0 |
|  | 13.1 | >99 (R) | 24.0 | 41.5 | >99 (R) | 24.0 |
|  | 91.6 | 51.4 (S) | 24.0 | 29.4 | 35.9 (S) | 24.0 |
|  | >99 | >99 (R) | 2.0 | >99 | 19.1 (R) | 0.5 |

Green highlight indicates the reduced bond on each compound.
<sup>a</sup> (1S,2R, 5S)-myrtanal as the major product. Double bond reduction results in altered CIP priority and hence inversion of assignment of this position.
<sup>b</sup> (2S,5S)-dihydrocarvone as the major product (*trans* configuration)
<sup>c</sup> (2S,5R)-dihydrocarvone as the major product (*cis* configuration)

**Figure 3:**
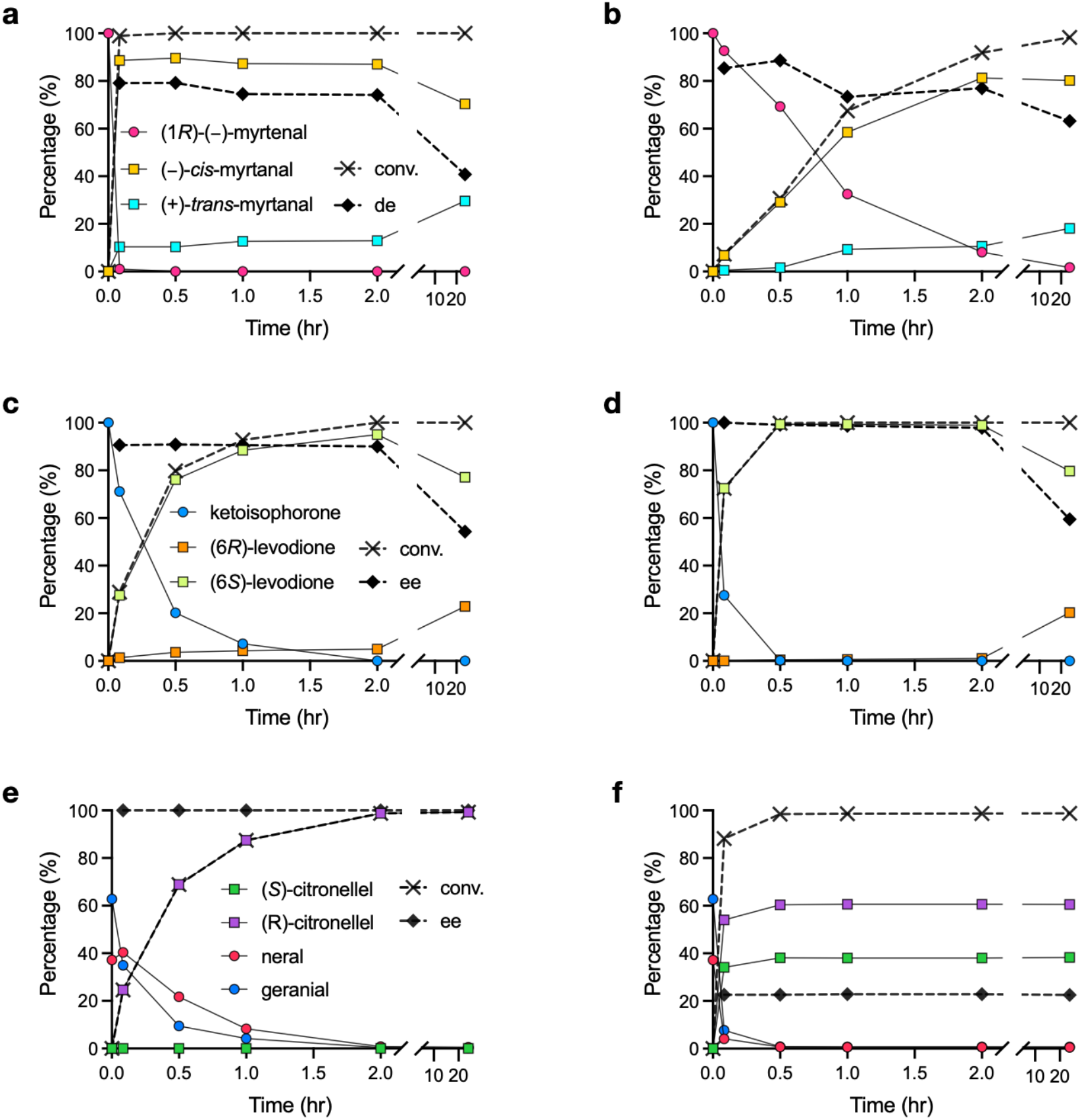
Time course of citral biotransformation by FDORs. Time course of the biotransformation with (1*R*)-(−)-myrtenal (**1a**), ketoisophorone (**2a**), and citral (**12a**) by FDORs. Relative concentrations of substrates and products, conversion, and de/ee are represented. Reactions were conducted in 50 mM Tris-HCl pH 8.0 containing 1.6 mM substrates, 73 µM F_420_, 2.2 µM FGD, 4.8 mM glucose-6-phohsphate, 2.9 µM FDOR and conducted at 30 °C. Final methanol concentration was 1.6% (v/v). Reactions were analyzed by chiral GC. Panel **a**, MSMEG_2027 with **1a**; panel **b**, MSMEG_2850 with **1a**; panel **c**, MSMEG_2027 with **2a**; panel **d**, MSMEG_2850 with **2a**; panel **e**, MSMEG_2027 with **12a**; panel **f**, MSMEG_2850 with **12a**.

(*S*)-(+)-Carvone **4a** was converted more slowly than (*R*)-(−)-carvone **5a**, however reduction of **4a** proceeded with greater stereoselectivity (94.8% and 78.8% *de* with MSMEG_2027 and MSMEG_2850, respectively) compared to **5a** (75.6 and 56.1% *de* with MSMEG_2027 and MSMEG_2850, respectively). As with both enantiomers of carvone, the 2-methylcyclohexenone **8a** and 2-methylcyclopenteneone **11a** were preferentially reduced to the (*S*)-configured products (**Table 3**), which suggests that these substrates bind in the same orientation within the active site. Substrates bearing a methyl group at the 3-position gave exclusively the (*R*)-configured products (**Table 3**) and thus the methyl groups are oriented on opposite faces of the ring compared to the products of the 2-methyl derivatives.

Citral **12a** (a 3:2 mixture of *trans*-**12a** and *cis*-**12a**) showed the greatest difference in stereoselectivity between the two enzymes, with 99% and 19.1% *ee* for MSMEG_2027 and MSMEG_2850, respectively. The time-course chromatogram of **12a** shows MSMEG_2027 accepted both isomers to yield almost exclusively (*R*)-**12b** after 120 minutes. In contrast, MSMEG_2850 showed complete conversion after 30 minutes but with a composition of 59.55% of (*R*)-**12b** and 40.45% of (*S*)-**12b** (**Figure 3**). This is remarkably close to the 3:2 ratio of geranial to neral in the citral sample used, which prompted us to hypothesise that MSMEG_2850 may convert neral (*Z*)-**12a** and geranial (*E*)-**12a** into (*S*)-**12b** and (*R*)-**12b**, respectively. Separate isomers of **12a** were obtained through Dess–Martin oxidation from geraniol and nerol (**Figure S4**). The enantioselectivity of reductions for (*E)*- and (*Z*)*-***12a** were determined by chiral GC (**Figure 3; Figure S5**). For each substrate, the *ee* of MSMEG_2027 exceeded 99% (*R*), as expected from that for **12a** (> 99% (*R*), **Table 3**). MSMEG_2850 converted (*E*)-**12a** to (*S*)-**12b** (*ee*, 53.9%) and (*Z*)-**12a** to (*R*)-**12b** (*ee*, >99%). This result refutes our initial hypothesis that MSMEG_2850 converts neral and geranial *exclusively* into (*S*)-**12b** and (*R*)-**12b**, respectively, however it does show that each isomer of **12a** is preferentially converted into a different enantiomer of **12b**.^52, 53^The interconversion of isomers of **12a** and **12b** is increased under alkaline conditions^53^ however, the results in **Figure 3** indicate that racemization of **12b** was negligible under the conditions tested here as *ee* did not decrease over time. It is noteworthy that of the tested substrates, geranial is the only one to exhibit opposite stereoselectivity between MSMEG_2027 and MSMEG_2850. Citral is particularly noteworthy for industrial catalysis as both FDORs preferentially produce the more desirable *(R)*-enantiomer, although MSMEG_2850 does so only with neral.^52, 53^

### Structure determination

To gain insight into the basis of the stereoselective reduction, we obtained the structure of full-length MSMEG_2850 at 1.36 Å. MSMEG_2850 crystallized in the cubic *I*4_1_32 space group with cell lengths of 128.8 Å (**Table 4; Figure S6**). A single polypeptide chain was contained in the asymmetric unit. MSMEG_2850 adopts the split β-barrel fold of FDOR-As with a core 7-stranded sheet interspersed with 4 helices (**Figure 4**). All but the first five residues could be built into the electron density. QtPISA analysis suggested that one of the crystal contacts represents a biological interface that results in formation of a dimer.^54^ This is supported by gel filtration data showing peaks corresponding to monomeric and dimeric species (**Figure S7**).

**Table 4:** Data collection, processing, and refinement statistics

|  |  |
| --- | --- |
| **PDB-ID** | 8D4W |
| **Data collection** |  |
| Space group | $I4_132$ |
| Cell dimensions |  |
| $a, b, c$ (Å) | 128.809, 128.809, 128.809 |
| $\alpha, \beta, \gamma$ (°) | 90, 90, 90 |
| Resolution (Å) | 45.54–1.35 (1.37–1.35)<sup>a</sup> |
| $R_{\text{merge}}$ | 0.135 (10.196)<sup>a</sup> |
| $R_{\text{pim}}$ | 0.016 (1.150)<sup>a</sup> |
| $I/\sigma I$ | 23.8 (0.8)<sup>a</sup> |
| $CC_{1/2}$ | 1.000 (0.350)<sup>a</sup> |
| Completeness (%) | 100 (100)<sup>a</sup> |
| Redundancy | 76.5 (79.0)<sup>a</sup> |
| **Refinement** |  |
| Resolution (Å) | 45.54–1.36 |
| No. reflections | 40068 (3947)<sup>a</sup> |
| $R_{\text{work}} / R_{\text{free}}$ | 18.8/20.2 |
| No. atoms |  |
| Protein | 1112 |
| Ligand/ion | 25 |
| Water | 132 |
| $B$-factors (Å<sup>2</sup>) | |
| Protein | 30.89 |
| Ligand/ion | 36.25 |
| Water | 38.59 |
| R.m.s. deviations |  |
| Bond lengths (Å) | 0.007 |
| Bond angles (°) | 0.99 |
| Ramachandran statistics |  |
| Favoured (%) | 98.5 |
| Allowed (%) | 1.5 |
| Outliers (%) | 0.0 |

<sup>a</sup> Values in parentheses are for the highest resolution shell

**Figure 4:**
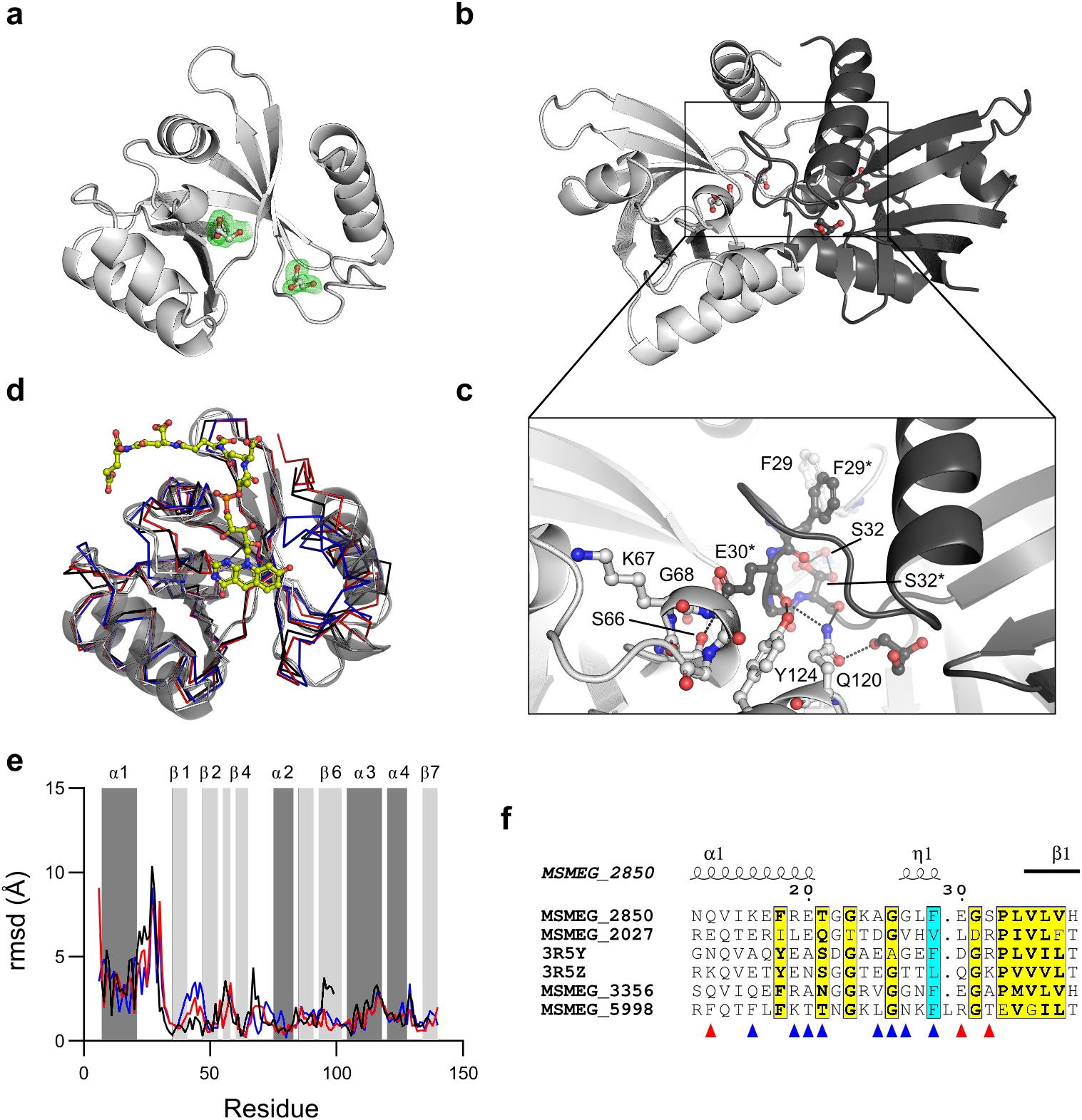
Structure of MSMEG_2850. **a** Cartoon representation of the asymmetric unit. Polder omit maps around glycerol molecules are contoured at 3σ **b** Cartoon representation of the dimer generated through application of the (*y* + ¼, *x* − ¼, −*z* −¼) symmetry operation. **C** Interchain contacts in the dimer interface. **D** Alignment of the MSMEG_2850 structure to other structures of full-length FDOR-A1s. MSMEG_2027 (6WTA), black; nfa33440 (3R5Y), red; nfa18080 (3R5Z), blue. For clarity only the F_420_-4 molecule of 6WTA and the cartoon of MSMEG_2850 are shown. **e** Per-residue rmsd for Cα atoms for the structure alignment. Coloring is as in panel **d. f** Portion of sequence alignment of FDOR-A1s showing that F29 aligns with other hydrophobic residues that interact with F_420_ (cyan). See **Figure S8** for full alignment.

The dimer interface consists of 12 interchain hydrogen bonds in addition to hydrophobic contacts. These occur between the α1 helices and between the loop regions between α1 and β1 and between β4 and α2, both of which contain a short 3_10_-helix (**Figure 4c**). E30* forms hydrogen bonds with the oxyanion hole formed by S66, K67 and G68, as well as Q120 and Y124. S32* also hydrogen bonds with Q120. There is likewise a hydrophobic interaction between F29 and F29*. Despite forming a dimer, the structure of MSMEG_2850 aligned to the monomeric structure of MSMEG_2027 in complex with F_420_ (PDB 6WTA) with a rmsd of 1.46 Å over 106 Cα atoms. Major differences in the backbone positions were observed in the loop region connecting α1 and β1 that forms part of the dimer interface (**Figure 4d** and **e**). In the structures of other full-length FDOR-As in complex with F_420_ this loop is contracted and contains a hydrophobic interaction between a sidechain and the phenolic ring of F_420_ (V30 in MSMEG_2027, F35 in 3R5Y, L33 in 3R5Z). Multiple sequence alignment suggests that F29 is the equivalent position in MSMEG_2850 (**Figure 4f**) which is positionally displaced to form part of the dimer interface.

This structure contrasts with that of dimeric MSMEG_2027 in which the N-terminal helices are domain-swapped and occupy the polyglutamate binding sites (PDB: 6XRI). In the MSMEG_2850 structure the helices do not domain swap, however domain-swapping of the loop region following the helices occupies the substrate and deazaflavin binding sites, probably stabilizing the apo-enzyme. The relative position and orientation of chains in the MSMEG_2027 and MSMEG_2850 dimers differ from each other and the two are not superimposable (**Figure S9**).

Two glycerol molecules originating from the cryobuffer could be built into the model of MSMEG_2850 (Figure 4a). The first of these is located at the ends of β2 and β4 in a pocket composed of the β2–β3 and β4–α2 loops, Q114, F121, and the α1–β1 loop of the opposite chain. This glycerol molecule makes polar contacts with both the amide NH and carbonyl oxygen of F64 and V73 as well as an ordered water molecule (**Figure S10**). In structures of homologous proteins with F_420_ bound this pocket is occupied by the pyrimidine ring of F_420_ and there is a conserved polar contact between N3 of the deazaflavin ring and the carbonyl of the equivalent residue of F64 (V65 in MSMEG_2027, V70 in 3R5Y, V68 in 3R5Z, see **Figures S11**,**12**). The second glycerol is in a pocket formed by the α1–β1 loop, the turn between β5 and β6, and the start of α4 of the opposite chain. Polar contacts occur between the amide NH and carbonyl of L34, the amide of G91, the carbonyl of G23 and the sidechain of Q120* (**Figure S10**). In contrast to the other ligand, this site is not associated with the binding of F_420_ or substrates. Polder omit maps^55^ show unambiguous density corresponding to both ligands (**Figure 4** and **Figure S10**).

Numerous attempts to obtain a structure of MSMEG_2850 in complex with F_420_ were unsuccessful, with the gel-purified complex dissociating within hours. Thermal denaturation curves in the presence of varying amounts of F_420_ suggest that the affinity of MSMEG_2027 and MSMEG_2850 are comparable under the conditions tested, with *T*_M50_ increasing by 9.1 ± 0.2 and 8.1 ± 0.2 degrees in the presence of a 10-fold excess of F_420_ compared to no F_420_ for MSMEG_2027 and MSMEG_2850, respectively (**Figure 5**). It may be that at the high concentrations required for crystallization, formation of the apo-dimer is more thermodynamically favourable than the monomeric holocomplex with F_420_.

**Figure 5:**
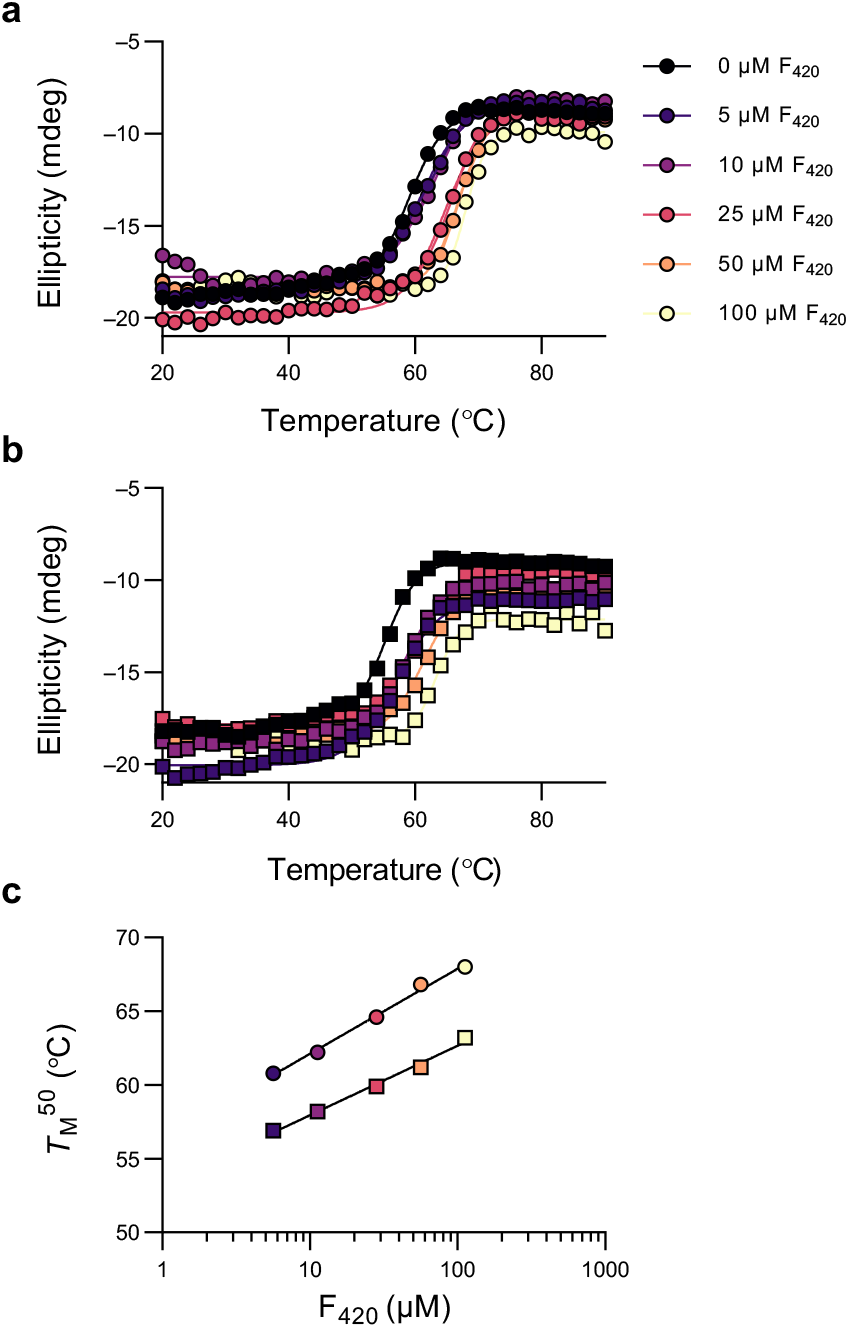
Effect of F_420_ concentration on melting temperature. Samples contained 10 µM FDOR with 0–100 µM F_420_ and were prepared in 10 mM sodium phosphate pH 7.5. Denaturation of **a** MSMEG_2027 and **b** MSMEG_2850 was followed at 218 nm. **c** *T*_*M*_^50^ increased linearly with log_10_[F_420_] over the concentration range tested with slopes of 5.8 ± 0.3 and 4.7 ± 0.3 °C, respectively for MSMEG_2027 (circles) and MSMEG_2850 (squares). In the absence of F_420_ the *T*_*M*_^50^ for MSMEG_2027 and MSMEG_2850 was 58.9 ± 0.2 and 55.1 ± 0.2 °C, respectively.

### Computational substrate docking

To rationalize the stereochemical outcomes observed in the enzymatic reductions we docked substrates into models of FDORs with F_420_H_2_ bound. A crystal structure of MSMEG_2027 was used (PDB 6WTA), while for MSMEG_2850, F_420_H_2_ was docked into the crystal structure of apo-MSMEG_2850 (**Figure S11**). Close inspection of the docked model of MSMEG_2850:F_420_H_2_ and comparison to the MSMEG_2027:F_420_H_2_ holo-enzyme structure showed the key interactions will fully conserved (**Figure S12**). Docking of the substrates into these holo-enzyme models was then performed (**Figure 6, Figure S13**). Poses were rejected if the β-carbon of the conjugated alkene was greater than 5 Å from the hydride of C5 of F_420_H_2_. In all cases an oxygen of the electron-withdrawing group hydrogen-bonds to the amide NH of G69, and additional hydrogen bonds to the hydroxyl of S67 as well as the amide NH of K68 were also observed in many poses, forming an oxyanion hole (numbering for MSMEG_2027). This SKGG motif is highly conserved amongst the FDOR-As,^10^ and the role of the serine hydroxyl as a hydrogen bond donor is consistent with MD simulations of Ddn with pretomanid.^21, 56^ For the prochiral substrates, the poses suggest a mechanism of hydride transfer to the β-carbon of the activated alkene followed protonation on the opposite face of the double bond. This results in net *trans*-addition of H_2_ across the double bond, consistent with the predominant mechanism of OYEs.^50^ The highly conserved tyrosine on helix 4 (Y126 in MSMEG_2027, Y124 in MSMEG_2850) could potentially serve as the proton donor, however mutagenesis and computational studies of Ddn suggest that the proton is derived from solvent.^21, 22^

**Figure 6.**
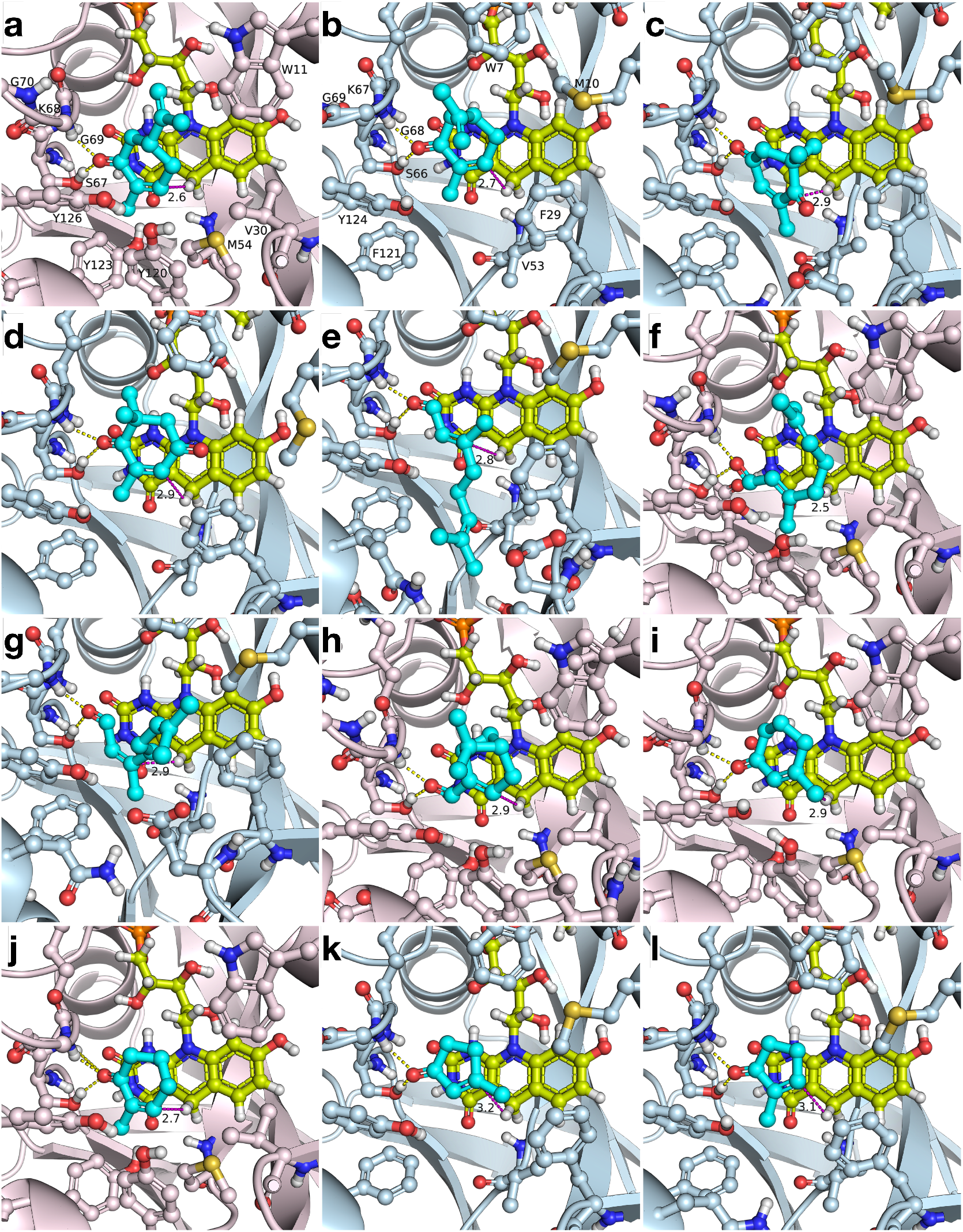
Representative induced-fit docking poses of prochiral substrates in MSMEG_2027 (pink) and _2850 (blue). Panel **a**, (*R*)-carvone **5a.**Panel **b**, (*S*)-carvone **4a**. Substrate **2a** bound in the C4-binding mode **c** and in the C1-binding mode **d**, both result in the same product. Isomers of citral **12a, e** and **f** geranial, which result in the opposite enantiomer of **12b**, and **g** neral. Panel **h, I, j, k** and **l** are **1a, 7a, 8a, 10a** and **11a**, respectively. Hydrogen bonds are shown as yellow dashed lines. The distance between the hydrogen of C5 in F_420_ and β-carbon of substrates aligned for hydride

The docking mode of *R*-carvone **5a** with MSMEG_2027 shows the isopropenyl group in an equatorial position, consistent with this being the lowest energy conformation^57^ (**Figure 6a**). Binding of (*S*)-carvone **4a** in the same conformation would cause a steric clash with a conserved tryptophan reside (W11 in MSMEG_2027, W7 in MSMEG_2850), suggesting that productive binding of **4a** may only occur in the less favorable axial conformation. This may partially explain the preference of **5a** and a substrate over **4a** by both enzymes, although flipping of this sidechain cannot be excluded as a possibility. In the case of ketoisophorone **2a**, two carbonyls are present in the substrate that could bind in the oxyanion hole. We obtained poses of both possible binding modes. In the first mode (**Figure 6b**), the C4-carbonyl binds in the oxyanion hole and the hydride of F_420_H_2_ is transferred to C2 on the *si*-face of the double bond. In the second mode, the C1-carbonyl binds in the oxyanion hole and the hydride is transferred to C3 on the *re*-face of the double bond (**Figure 6c**). Both poses thus give rise to the observed (*S*)-product assuming net *trans*-addition.

We obtained different binding modes for *trans*-**12a** with MSMEG_2027 and MSMEG_2850, with the *si*- and *re*-faces, respectively, being presented to the cofactor (**Figure 6**). Assuming net trans-addition of H_2_ in each case, these would result in (*R*)- and (*S*)-**12b**, respectively. In MSMEG_2850, the isobutylene tail of *trans*-**12a** can be placed downwards interacting with loop 1 (between α1 and β1) and loop 10 (between α3 and α4) due to the substitution of less bulky residues A118 and F121 (**Figure 6**), compared to corresponding Y120 and Y123 of MSMEG_2027. In addition, the more closed conformation of α1 towards loop 5 (between β4 and α2) in the F_420_ cleft of MSMEG_2850 than that of MSMEG_2027 (**Figure 6**) prevents the isobutylene tail of *trans*-**12a** from being oriented upwards as in **Figure 6**, leading to reversed binding modes between MSMEG_2027 and MSMEG_2850. Neral was bound with its *re*-face oriented towards F_420_H_2_ in both enzymes, generating (*R*)-**12b**. However, the placement of the isobutylene tail was distinct: while MSMEG_2027 aligned it upwards along the ribityl chain of F_420_, MSMEG_2850 placed it over the xylene ring (A) of F_420_ (**Figures 6 and S13**). These suggest that rather than neral and geranial being exclusively reduced into a single enantiomer of citronellal, two binding modes are possible for each substrate that each result in a different enantiomer. This has been shown to occur with OYE2y-mediated reduction of citral^53, 58^ as well as with other FDORs.^7^

For (*R*)-myrtenal **1a**, the geminal dimethyl bridgehead points away from F_420_ (**Figure 6**). The obtained poses would yield *trans*-**1b** following net *trans*-addition across the double bond, however the *cis*-configured product is the major product obtained with both enzymes. This implies that for this substrate net *cis* addition of H_2_ across the double bond is the major pathway. It is unclear why this substrate does not follow the general trend. No literature data is available for the enantioselectivity for the reduction of **1a** by OYE family ene reductases for comparison, although divergent mechanisms for the reduction of (*R*)- and (*S*)-perillaldehyde by the non-flavin double bond reductase Ltb4dh has been shown.^59^

Poses of the endocyclic enones **7a, 8a, 10a** and **11a** (**Figure 6i, j, k**, and **I**, respectively), are like that of **4a**. Addition of the hydride to C3 results in the observed (*R*)-configuration in the products, while protonation on the opposite face results in the observed *S*-configuration of **8b** and **11b** (**Scheme 1**). Both enzymes transformed **11a** with substantially lower stereoselectivity than **8a** (91.9% vs 51.4%, and 72.0% vs 35.9%, **Table 3**). One possible explanation is that the smaller **11a** may be better able to bind in a catalytically productive flipped orientation compared to **8a**, although no difference in selectivity was observed for the 3-methyl products **7b** and **10b**. Another possibility is that while hydride addition is selective for one face only, protonation of the proximal face represents a competing pathway and the smaller size of **11a** makes net *cis*-hydrogenation more favorable for 5-membered rings compared with 6-membered rings. **11b** may also have a greater propensity to racemize under the assay conditions compared to **8b**. Many OYEs have been shown to have much poorer enantioselectivity with **11a** than with **8a**.^60–62^

**Scheme 1:**
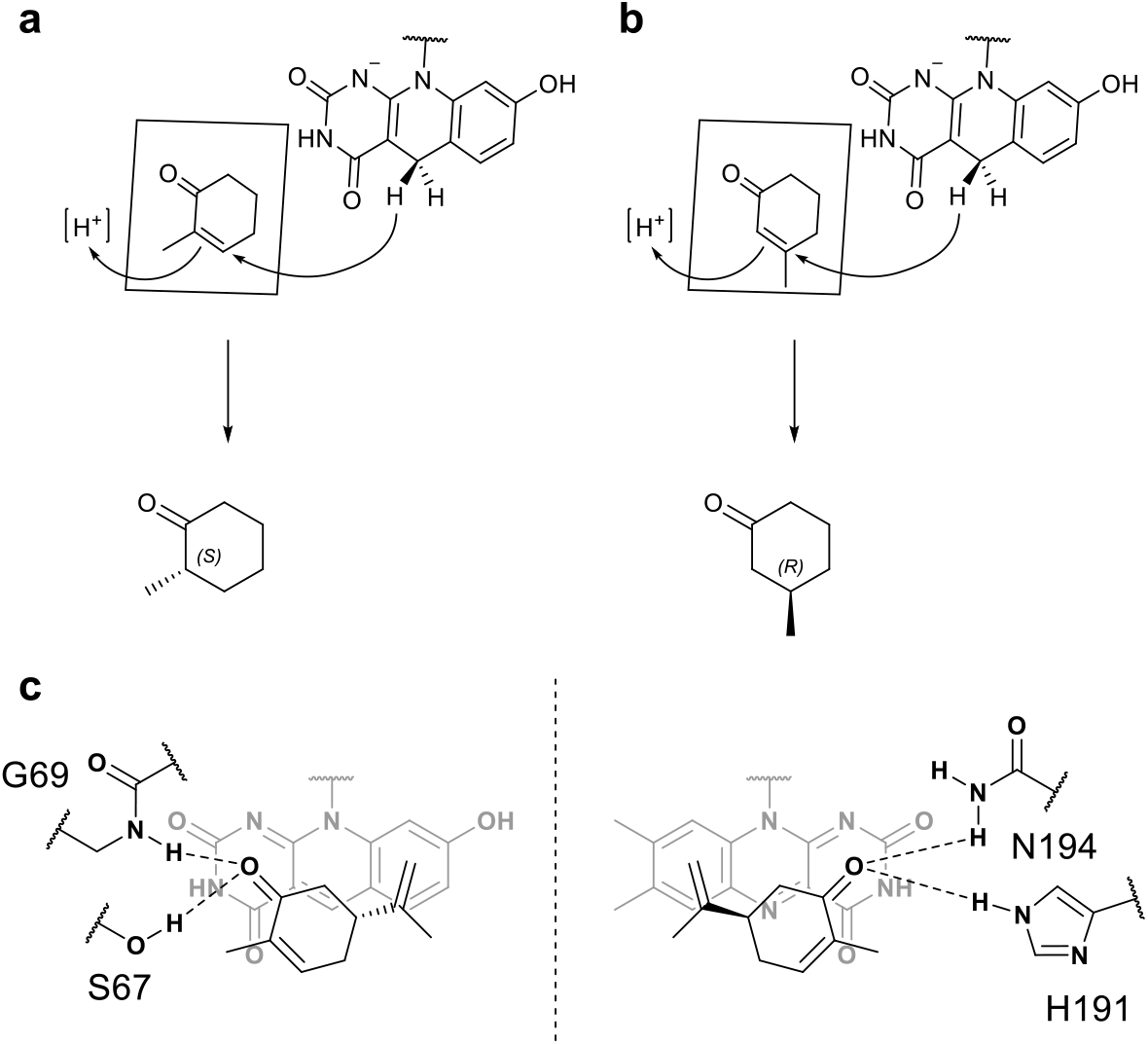
Proposed mechanism of action for F_420_-mediated reductions. The observed stereochemistry of products **8b** (a) and **7b** (b) can be rationalized by transfer of a hydride from F_420_H_2_ to the proximal side of the β-carbon followed by protonation of the α-carbon on the distal side. In FDORs the *re*-face of the deazaflavin is presented to the substrate and the oxyanion hole is located on the left as draws. In the OYE family the *si*-face of the flavin is presented to the substrate and the oxyanion hole is on the right as drawn, thus mirroring the orientation of the FDORs through the plane indicated by the dashed line. Residue numbering is for MSMEG_2027 and OYE1, respectively.

### Impact of electron-withdrawing group strength on activity

There have been several attempts to correlate ene-reductase activity with measures of the intrinsic reactivity of the substrates. Romano *et al*. used density functional theory (DFT) to calculate the electrostatic potential of the carbon undergoing hydride transfer for a series of cyclohexanone derivatives.^63^ Freund *et al*. used calculated hydride affinities combined with docking scores to assign compounds as probable substrates of a family of enoyl acyl carrier protein reductases.^64^ In both cases, the authors found that enzyme activity did not strongly correlate with the intrinsic reactivity of the substate, but rather that the steric effects of substituents on binding within the active site played the major role in determining reaction rate. However, the substrates tested did not cover a wide range of EWG groups. The Hammett equation is widely used to predict relative reaction rates for reagents baring substituents with varying electron donating and withdrawing character.^65^ We compared the log of the specific activity of compounds **1 3a**–**17a**, which differ only by their EWG, with the intrinsic electronic component of the Hammett σ_p_, *σ*_e_(*ω*), calculated by Domingo et. al., for a series of substituted ethylenes.^66^

The results, shown in **Figure S14** on and **Table S4**, show that there is a positive correlation between specific activity and *σ*_e_(*ω*) for both MSMEG_2027 and MSMEG_2850. Interestingly, the activity of MSMEG_2850 with **17a** was greater than that of **16a** consistent with the *σ*_p_ value. *R*^2^ values for the fit were 0.584 for MSMEG_2027 and 0.704 for MSMEG_2850, respectively (**Table S4**). These results show that the strength of the EWG strongly affects activity for the functional groups tested. Importantly, each of the functional groups tested is isosteric and thus the influence of EWG geometry is minimized.

## Discussion

In this study, we took advantage of the fluorescence of F_420_ to rapidly screen multiple enzymes for activity against a panel of substrates, allowing us to rapidly identify homologues with broad substrate ranges and high activities using a single manually operated 96-well plate reader. This approach is more rapid and parallelizable than the GC-based approaches typically used to screen FMN-dependent ene-reductases and can be easily adapted for automated plate reader setups. Although colorimetric assays for these ene-reductases have been developed they require additional enzymes and dyes as spectral overlap between many substrates and NAD(P)H renders direct spectrophotometric quantitation problematic.^67^ This makes FDORs highly amenable to high-throughput screening strategies, which is particularly useful for directed evolution experiments.

In comparison to previous work, the activity and selectivity of MSMEG_2027 and MSMEG_2850 is broadly similar to the previously characterized FDOR-A1s FDR-Mha, FDR-Rh1 and FDR-Rh2.^7^ With **2a**, both MSMEG_2027 and MSMEG_2850 showed higher turnover compared to the previously characterized enzymes, with MSMEG_2850 showing both the highest activity and the highest *ee* of any FDOR characterized thus far (**Table 3**). With (*R*)-carvone **5a** MSMEG_2027 showed the highest *de* of any enzyme, although this was still a modest 75.6% (**Table 3**). Reduction of substrates bearing a methyl substituent on the β-carbon (**7a** and **10a**) gave exclusively the (*R*)-configured products. The outcome for the corresponding α-methyl compounds varied considerably between enzymes, with MSMEG_2027 and MSMEG_2850 favoring the (*S*)-configured product **8b** much more strongly than the previously characterized enzymes (91.9 and 72.0% *ee, cf*. 43% for FDR-Rh2). Likewise, for **11a** the (*S*)-configured product is more strongly favored by MSMEG_2027 and MSMEG_2850 than for the previously characterized enzymes.

This study also developed our understanding of the catalytic mechanism of these enzymes. Studies of the OYE family of ene-reductases have concluded that the basis of their selectivity is a result of several factors: (i) The carbonyl (or equivalent) of the electron-withdrawing group preferentially binds to (and developing negative charge along the reaction coordinate is stabilized by) the conserved active histidine and asparagine/histidine pair and that the β-carbon must be oriented above N5 in a way that facilitates hydride transfer; (ii) given these constraints there are two possible faces of the double bond that could be presented to the cofactor, differing by a 180 degree rotation (flipping) along the C1–C2–C3 axis; (iii) the reaction typically (but not exclusively) proceeds with hydride and proton addition occurring on opposite faces of the double bond; (iv) for a given substrate and mode of hydride transfer (cis-or trans) a single product is obtained, which is altered by a change to either the binding mode or the mode of hydride addition; (v) steric interactions between substrate substituents and active site residues determine the relative energy of each binding mode and thereby determine the selectivity of the reaction.^68–70^ In both the FDOR-As and OYEs the oxyanion hole into which substrate carbonyls bind is located above the pyrimidine ring of the (deaza)flavin. However, it is the *re*- and *si*-face, respectively, of the cofactor that is presented to the substrate in these families. Thus, the binding site is effectively mirrored in the FDORs relative to the OYEs. If the active sites favor the same relative orientation of the substrate and cofactor the “classical” binding mode of OYEs corresponds to the “flipped” mode of FDORs and *vice-versa* (**Scheme 1**) and therefore the two families will give generally opposite selectivity given that (ii), (iii) and hold. This appears to be the major determinant of the observed opposite selectivity between OYEs and FDORs,^61, 62^ as has been suggested previously.^7^ Indeed, this switch in selectivity is observed with *Ph*ENR and *Tt*ENR, which also present the *re*-face of the cofactor to the substrate.^43^ We hypothesize that this trend might generalize to other substrates for which OYEs show strong selectivity, barring any substantial deviations in active site geometry. If this holds true, then the FDORs present a possible alternative to mutagenesis in achieving the desired enantiomeric product. Indeed, this was recently used by Venturi *et al*., in their enantiodivergent synthesis of cyclic halohydrins containing three contiguous stereogenic centers.^71^

## Conclusion

Reduction of α,β-unsaturated compounds, including many classic OYE substrates, is widespread amongst the FDOR-As, with the FDOR-A1 subgroup showing particularly high activities and a broad substrate range. From this group we selected MSMEG_2027 and MSMEG_2850 for more detailed characterization. Apparent steady state kinetic parameters typically revealed a *k*_cat_ in the range of 0.1–1.0 s^−1^ and a *K*_M_ in the low millimolar to high nanomolar range, as is typical of non-native substrates. Structural characterization of MSMEG_2850 revealed a novel dimeric structure distinct both from the FDOR-Bs and from that of MSMEG_2027. Induced fit docking of substrates allowed the observed stereochemical outcomes to be mechanistically rationalized as net *trans*-addition of H_2_ across the alkene. While mechanistically analogous to the OYE family of ene-reductases, the active site geometry of each family is effectively a mirror reflection of the other. As a consequence, the preferred binding mode of substrates is flipped between the two families. We propose that this trend will generalize to other substrates and therefore that the FDOR-A1s are excellent candidates for expanding the toolbox of ene-reductases.

## Data availability

The coordinates for the structure of MSMEG_2850 has been deposited in the Protein Data Bank with accession code 8D4W. The UNIPROT IDs of proteins described in this paper are: RER_09240, C0ZR82; MSMEG_2850, A0QW82; MSMEG_2027; A0QU01, MSMEG_5998, A0R4Y6; RV3547, P9WP15; MSMEG_5215, A0R2S3; MSMEG_3909, A0QZ62.

## Acknowledgements

This research was undertaken in part using the MX2 beamline at the Australian Synchrotron, part of ANSTO, and made use of the Australian Cancer Research Foundation (ACRF) detector. This work was supported by the CSIRO SynBio FSP, the ARC Centre of Excellence in Synthetic Biology (CE200100029) and the ARC Centre of Excellence in Peptide and Protein Science (CE200100012).

