## Supplementary Information for "Asymmetric ene-reduction of α,β-unsaturated compounds by F_420_-dependent oxidoreductases A (FDOR-A) enzymes from *Mycobacterium smegmatis*"

**Table S1:** Plasmids used in this study

| Enzyme | Clade | Vector | Reference |
|---|---|---|---|
| MSMEG_2027 | A1 | pETMCSIII | Ahmed *et al.*[1] |
| MSMEG_2850 | A1 | pDEST17 | Taylor *et al*[2] |
| MSMEG_2850 | A1 | pETMCSIII | This study |
| MSMEG_3909 | A3 | pDEST17 | Lapalikar *et al.*[3] |
| MSMEG_5215 | A3 | pDEST17 | Lapalikar *et al*[3] |
| MSMEG_5998 | A1 | pDEST17 | Taylor *et al* |
| RER_09240 | A1 | pDEST17 | Lapalikar *et al*[3] |
| Rv3547 | A1 | pMal-c2x | Lee *et al.*[4] |
| FGD | — | pET14b | Taylor *et al*[2] |

**Table S2:** Retention times from achiral GC/MS analysis

| Entry | Substrate | $R_t$ (min) | Product(s) | $R_t$ (min) |
|---|---|---|---|---|
| 1 | **1a** | 19.76 | *trans*-**1b**<br>*cis*-**1b** | 19.49<br>19.63 |
| 2 | **2a** | 18.15 | **2b** | 18.57 |
| 3 | **3a** | 15.51 | **3b** | — |
| 4 | **4a** | 20.73 | *trans*-**4b** | 19.77 |
| 5 | **5a** | 20.74 | *trans*-**5b**<br>*cis*-**5b** | 19.78<br>19.93 |
| 6 | **6a** | 11.47 | **6b** | 10.36 |
| 7 | **7a** | 15.66 | **7b** | 12.18 |
| 8 | **8a** | 13.78 | **8b** | 12.16 |
| 9 | **9a** | 8.69 | **9b** | 7.74 |
| 10 | **10a** | 8.76 | **10b** | 5.90 |
| 11 | **11a** | 10.77 | **11b** | 8.98 |
| 12 | **12a** | 20.73 (*cis*)<br>21.28 (*trans*) | **12b** | 18.82 |
| 13 | **13a** | 13.55 | **13b** | 11.60 |
| 14 | **14a** | 11.91 | **14b** | 14.49 |
| 15 | **15a** | 17.49 | **15b** | 15.31 |
| 16 | **16a** | 17.32 | **16b** | — |
| 17 | **17a** | 18.34 | **17b** | — |
| 18 | **18a** | 21.14 | **18b** | 19.17 |
| 19 | **19a** | 21.06 | **19b** | 18.67 |
| 20 | **20a** | 18.76 | **20b** | 16.73 |

**Table S3:** Chiral GC methods and retention times

| Method | Substrates | Column[a] | Conditions | $R_t$ (min) |
|---|---|---|---|---|
| 1 | **1a** | 1 | 90 °C for 3 min; 1.5 °C min$^{-1}$ until 123 °C; hold for 5 min; 10 °C min$^{-1}$ until 180 °C; hold for 2 min | **1a**, 21.64<br>*cis*-**1b**, 21.85<br>*trans*-**1b**, 23.27 |
| | **4a** | | | **4a**, 26.19<br>*cis*-**4b**, 21.41<br>*trans*-**4b**, 22.36 |
| | **5a** | | | **5a**, 26.38<br>*cis*-**5b**, 21.34<br>*trans*-**5b**, 22.27 |
| 2 | **2a** | 2 | 90 °C for 2 min; 4 °C min$^{-1}$ until 115 °C; hold for 10 min; 10 °C min$^{-1}$ until 180 °C; hold for 2 min | **2a**, 11.85<br>(*R*)-**2b**, 12.36<br>(*S*)-**2b**, 12.45 |
| 4 | **7a** | 1 | 40 °C for 0 min; 5 °C min$^{-1}$ until 95 °C; hold for 10 min; 10 °C min$^{-1}$ until 150 °C; hold for 10 min | **7a**, 24.29<br>(*R*)-**7b**, 15.95<br>(*S*)-**7b**, 16.26 |
| 3 | **8a** | 2 | 80 °C for 6.5 min; 10 °C min$^{-1}$ until 130 °C; hold for 3 min | **8a**, 10.58<br>(*S*)-**8b**, 9.93<br>(*R*)-**8b**, 10.05 |
| 6 | **10a** | 1 | 40 °C for 0 min; 2 °C min$^{-1}$ until 55 °C; hold for 40 min; 10 °C min$^{-1}$ until 150 °C; hold for 5 mins | **10a**, 51.46<br>(*R*)-**10b**, 21.52<br>(*S*)-**10b,** 22.08 |
| 5 | **11a** | 1 | 40 °C for 0 min; 5 °C min$^{-1}$ until 68 °C; hold for 15 min; 10 °C min$^{-1}$ until 160 °C; hold for 2 mins | **11a**, 16.73<br>(*R*)-**11b**, 11.95<br>(S)-**11b**, 12.53 |
| 7 | **12a** | 1 | 70 °C for 2 min; 1 °C min$^{-1}$ until 90 °C; hold for 10 min; 4 °C min$^{-1}$ until 130 °C; hold for 2 mins; 10 °C min$^{-1}$ until 150 °C; hold for 1 min | *cis*-**12a**, 40.71<br>*trans*-**12a**, 43.24<br>(*S*)-**12b**, 30.16<br>(*R*)-**12b**, 30.32 |

[a] Column 1: CycloSil-B (30 m × 0.25 mm × 0.25 µm); Column 2: Chiral Dex CB (25 m × 0.32 mm × 0.25 µm).

**Table S4:** Comparison of relative velocities with EWG strength

| Substrate | $\sigma_e(\omega)$[a] | $k_{obs}$ ($s^{-1}$)[b] | |
|---|---|---|---|
| | | MSMEG_2027 | MSMEG_2850 |
| **13a** | 0.79 | 48.2 | 7.9 |
| **14a** | 0.57 | 42.9 | 5.4 |
| **15a** | 0.51 | 3.6 | 3.8 |
| **16a** | 0.50 | 1.5 | 0.5 |
| **17a** | 0.46 | 0.5 | 0.9 |
| $R^2$ | | 0.5846[c] | 0.7038[c] |
| Slope | | 2.166[c] | 1.756[c] |
| Y intercept | | −0.0078[c] | −0.4754[c] |

[a] From Domingo *et. al.*,[5]

[b] Data obtained from **Table 1** in main text.

[c] Semilog regression of $k_{obs}$ vs $\sigma_e(\omega)$.

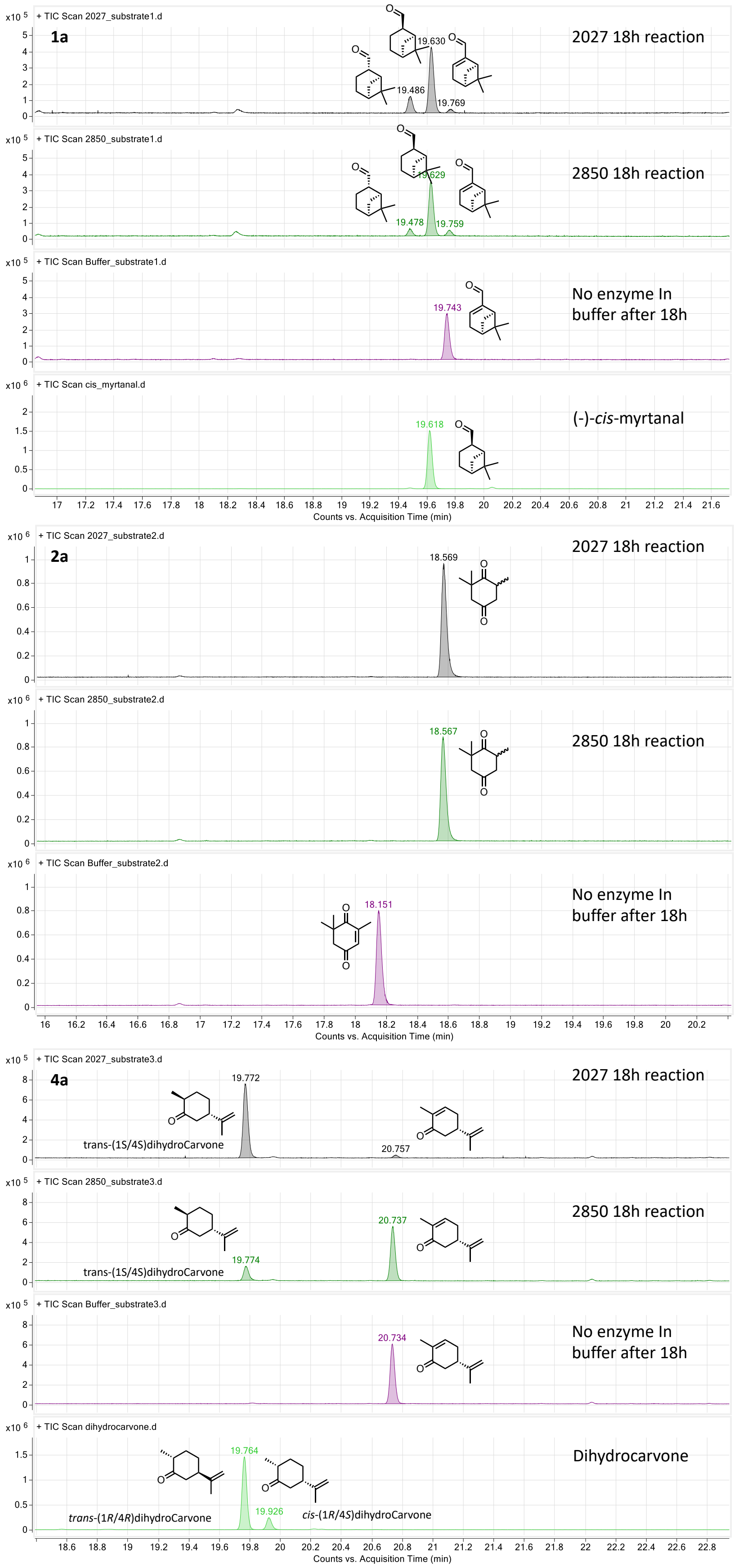


Figure S1. Achiral GC/MS of substrate conversion. Reactions contained 72.5 μM $F_{420}$, 2.9 μM FDOR, 1.6 mM substrate, 6.5 mM G6P and 1.4 μM Fgd in 100 μL volume

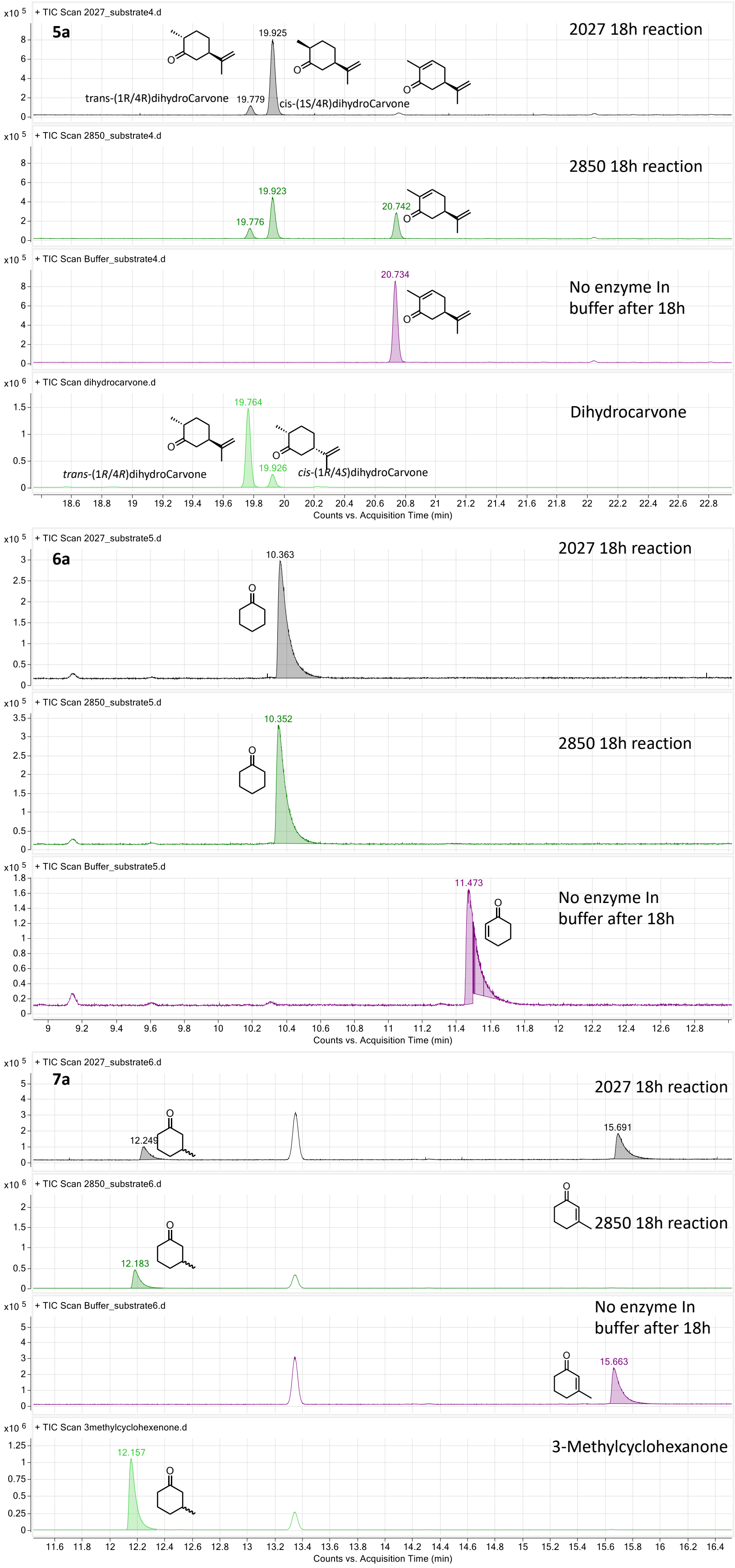


Figure S1 **(cont.):** Achiral GC/MS of substrate conversion. Reactions contained 72.5 µM $F_{420}$, 2.9 µM FDOR, 1.6 mM substrate, 6.5 mM G6P and 1.4 µM Fgd in 100 µL volume

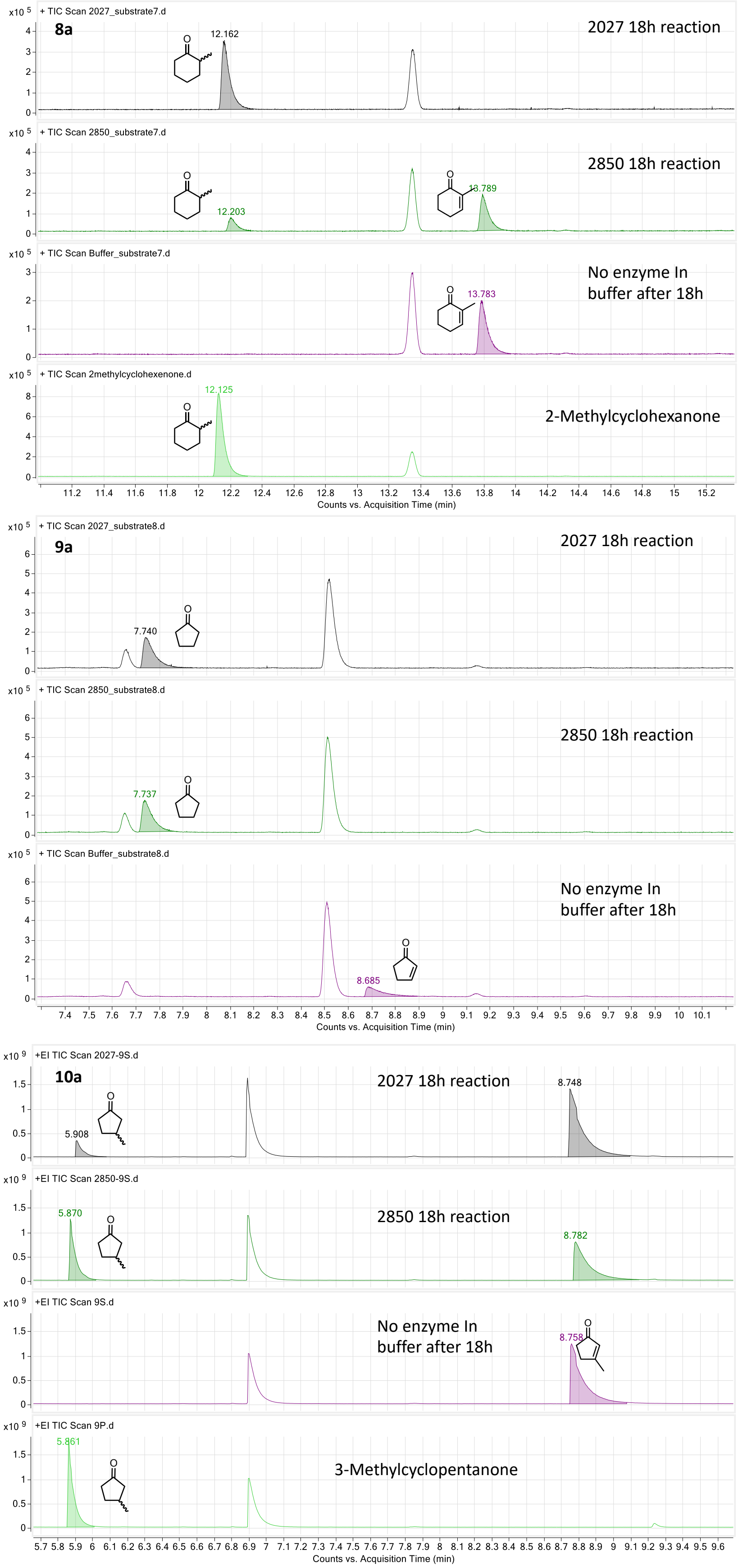


Figure S1 **(cont.):** Achiral GC/MS of substrate conversion. Reactions contained 72.5 μM $F_{420}$, 2.9 μM FDOR, 1.6 mM substrate, 6.5 mM G6P and 1.4 μM Fgd in 100 μL volume

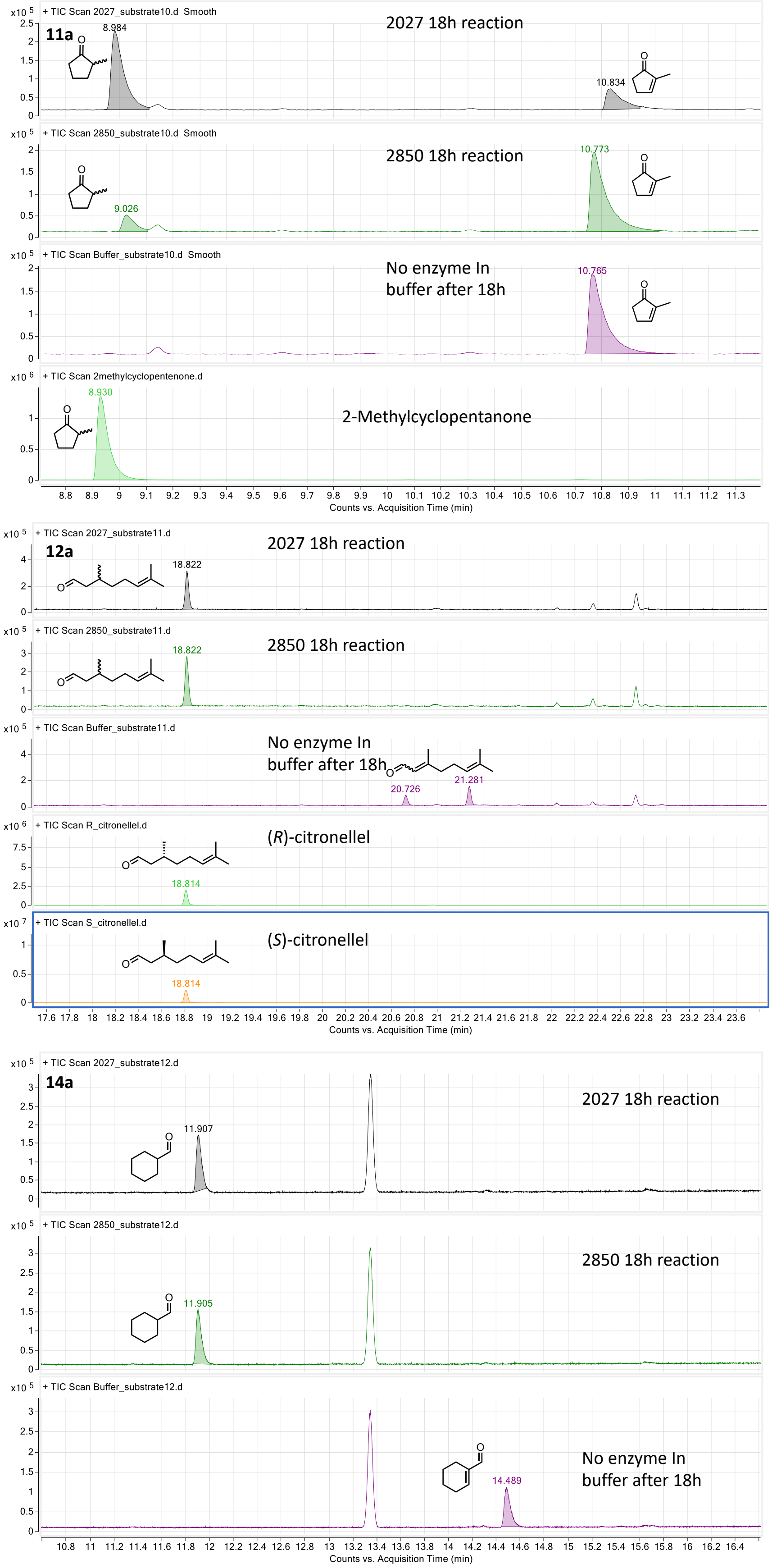


Figure S1 **(cont.):** Achiral GC/MS of substrate conversion. Reactions contained 72.5 μM $F_{420}$, 2.9 μM FDOR, 1.6 mM substrate, 6.5 mM G6P and 1.4 μM Fgd in 100 μL volume

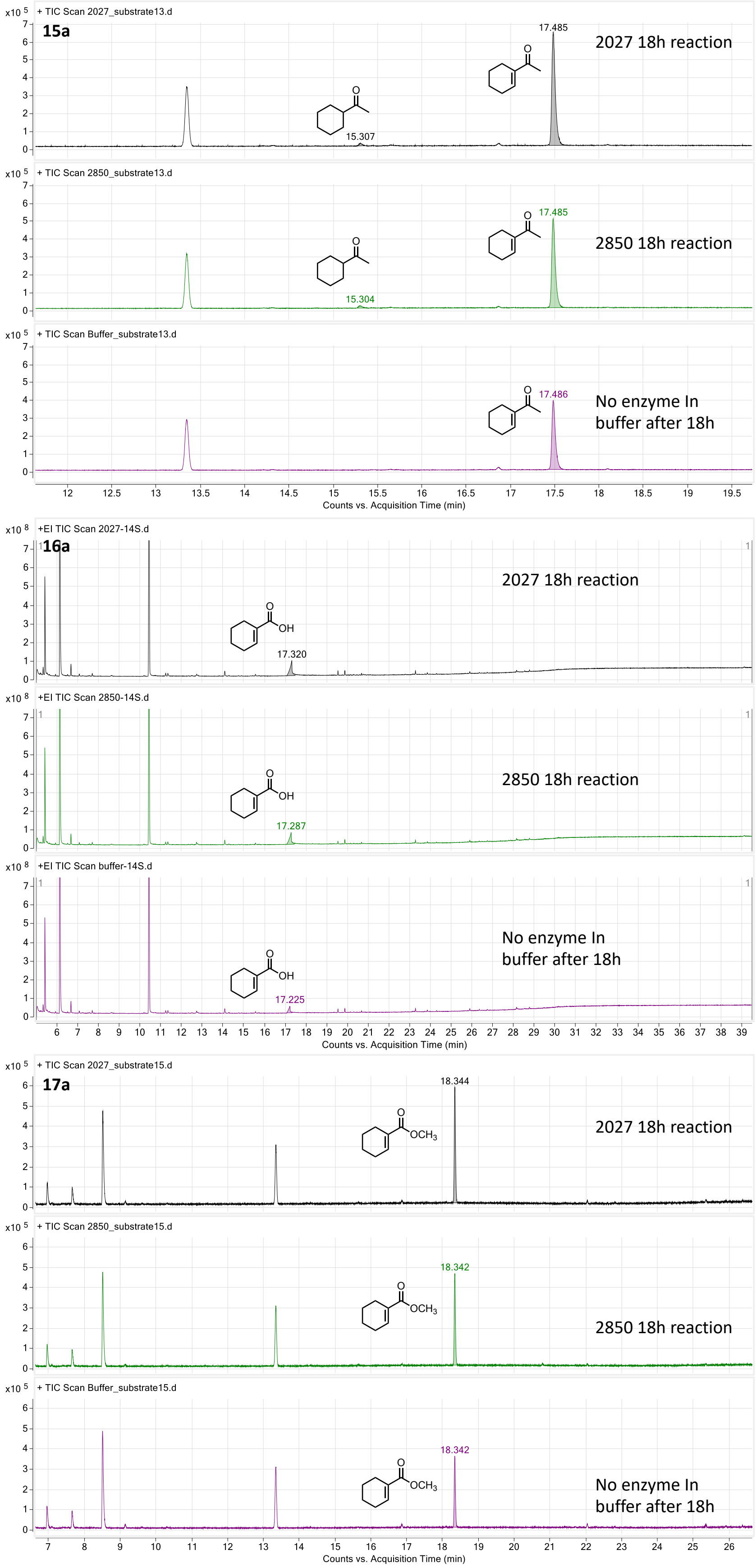


Figure S1 **(cont.):** Achiral GC/MS of substrate conversion. Reactions contained 72.5 µM $F_{420}$, 2.9 µM FDOR, 1.6 mM substrate, 6.5 mM G6P and 1.4 µM Fgd in 100 µL volume

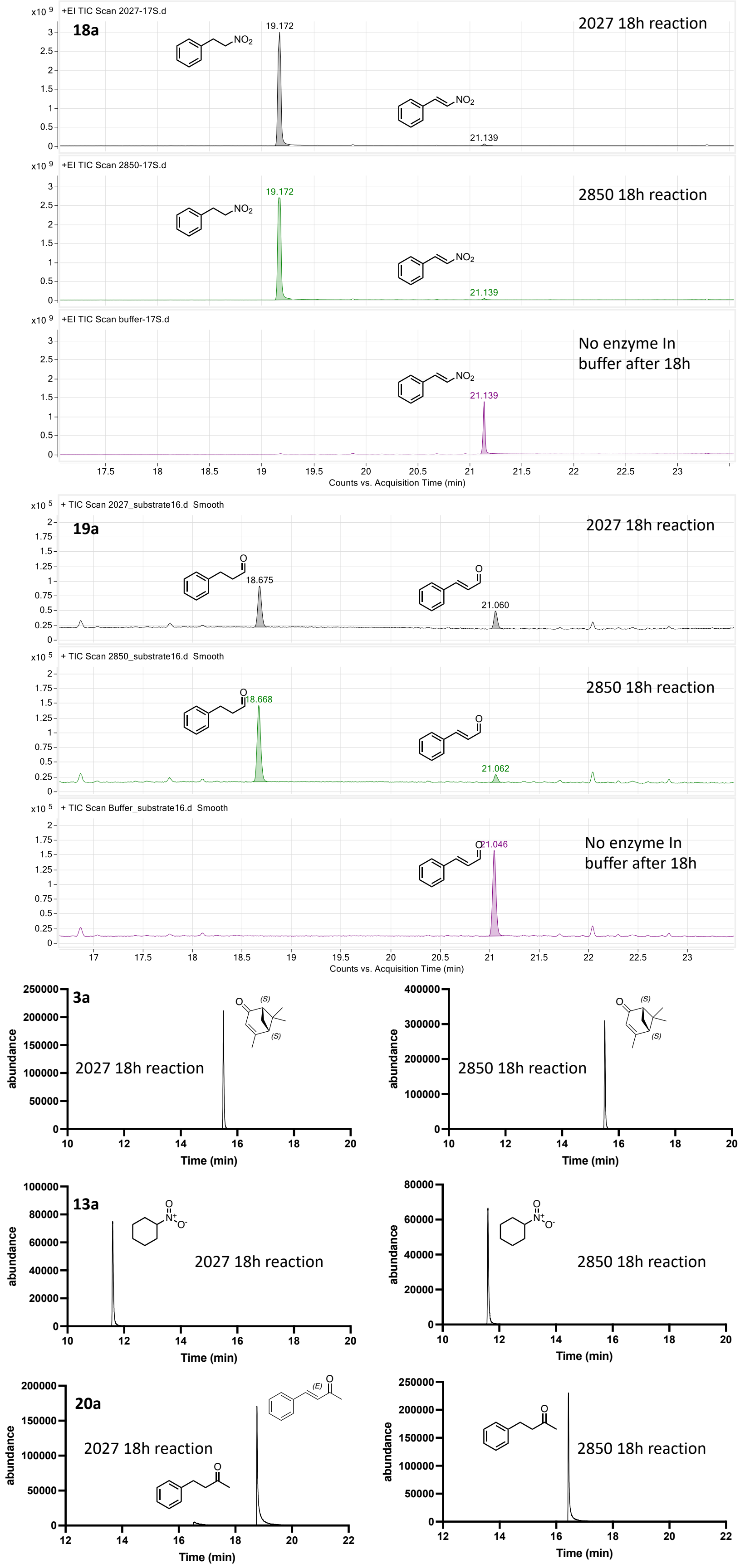


**Figure S1 (cont.):** Achiral GC/MS of substrate conversion. Reactions contained 72.5 µM $F_{420}$, 2.9 µM FDOR, 1.6 mM substrate, 6.5 mM G6P and 1.4 µM Fgd in 100 µL volume.

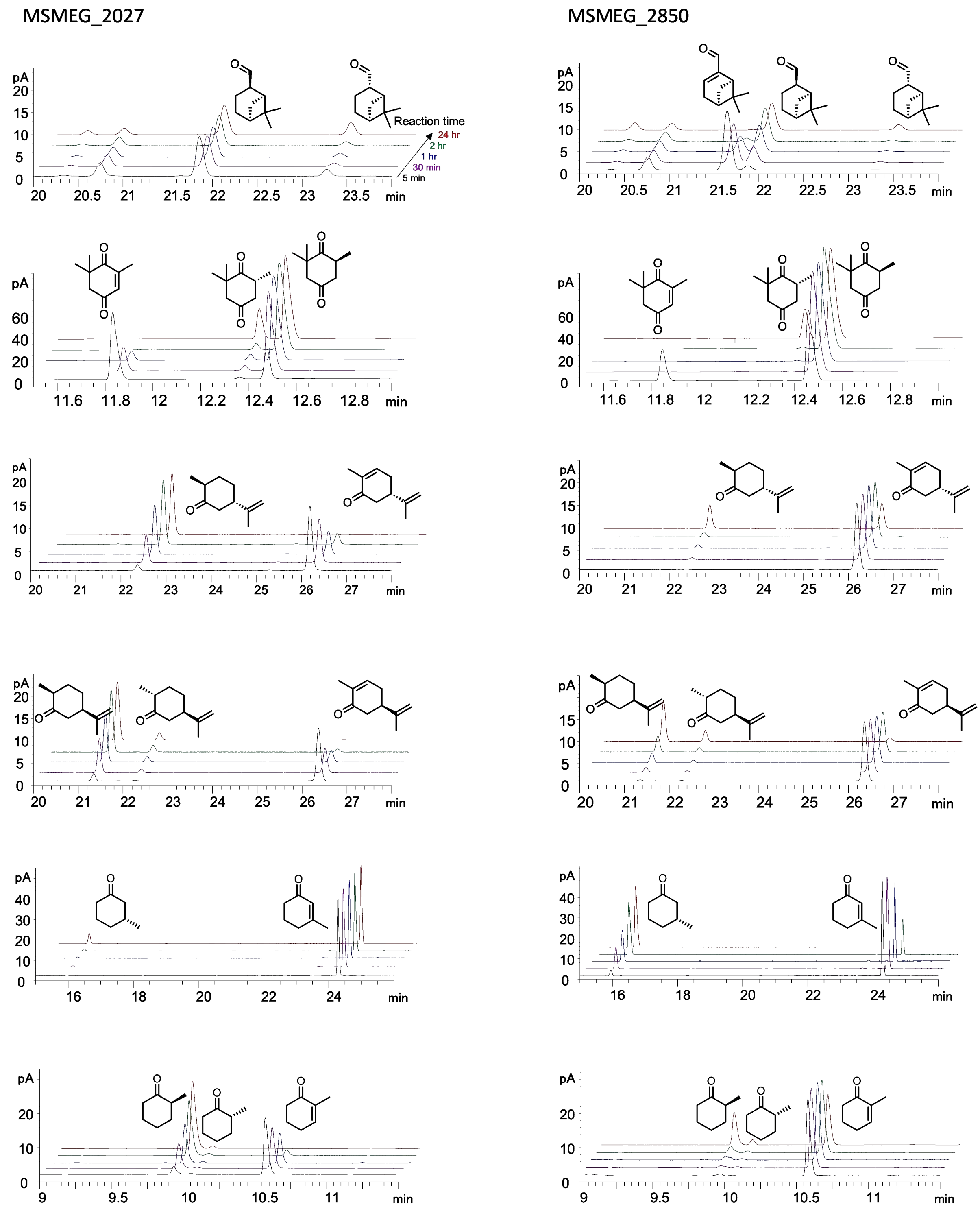


**Figure S2**. Time course chiral GC of reduction of prochiral substrates. Reactions contained 72.5 μM $F_{420}$, 2.9 μM FDOR, 1.6 mM substrate, 6.5 mM G6P and 1.4 μM Fgd in 775 μL volume. Samples were taken for analysis at 5 minutes, 30 minutes, 1 hour, 2 hours, and 24 hours

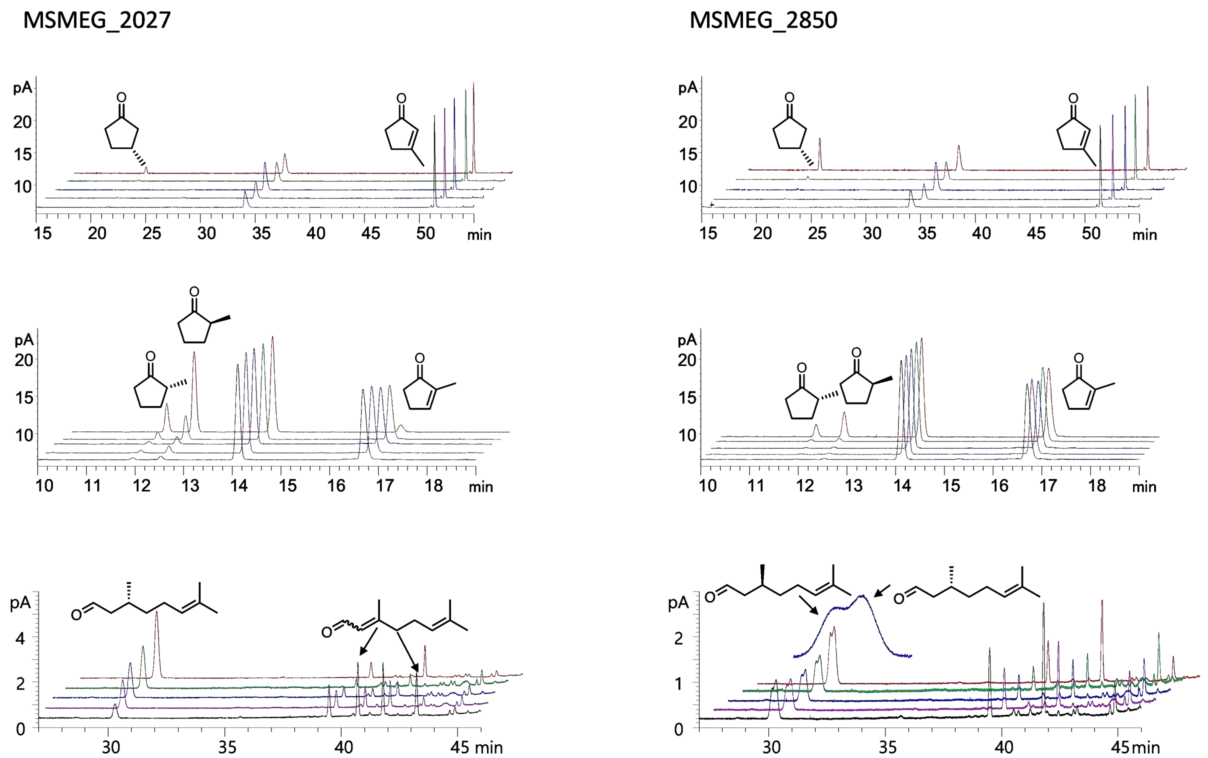


**Figure S2 (cont.):** Time course chiral GC of reduction of prochiral substrates. Reactions contained 72.5 µM $F_{420}$, 2.9 µM FDOR, 1.6 mM substrate, 6.5 mM G6P and 1.4 µM Fgd in 775 µL volume. Samples were taken for analysis at 5 minutes, 30 minutes, 1 hour, 2 hours, and 24 hours

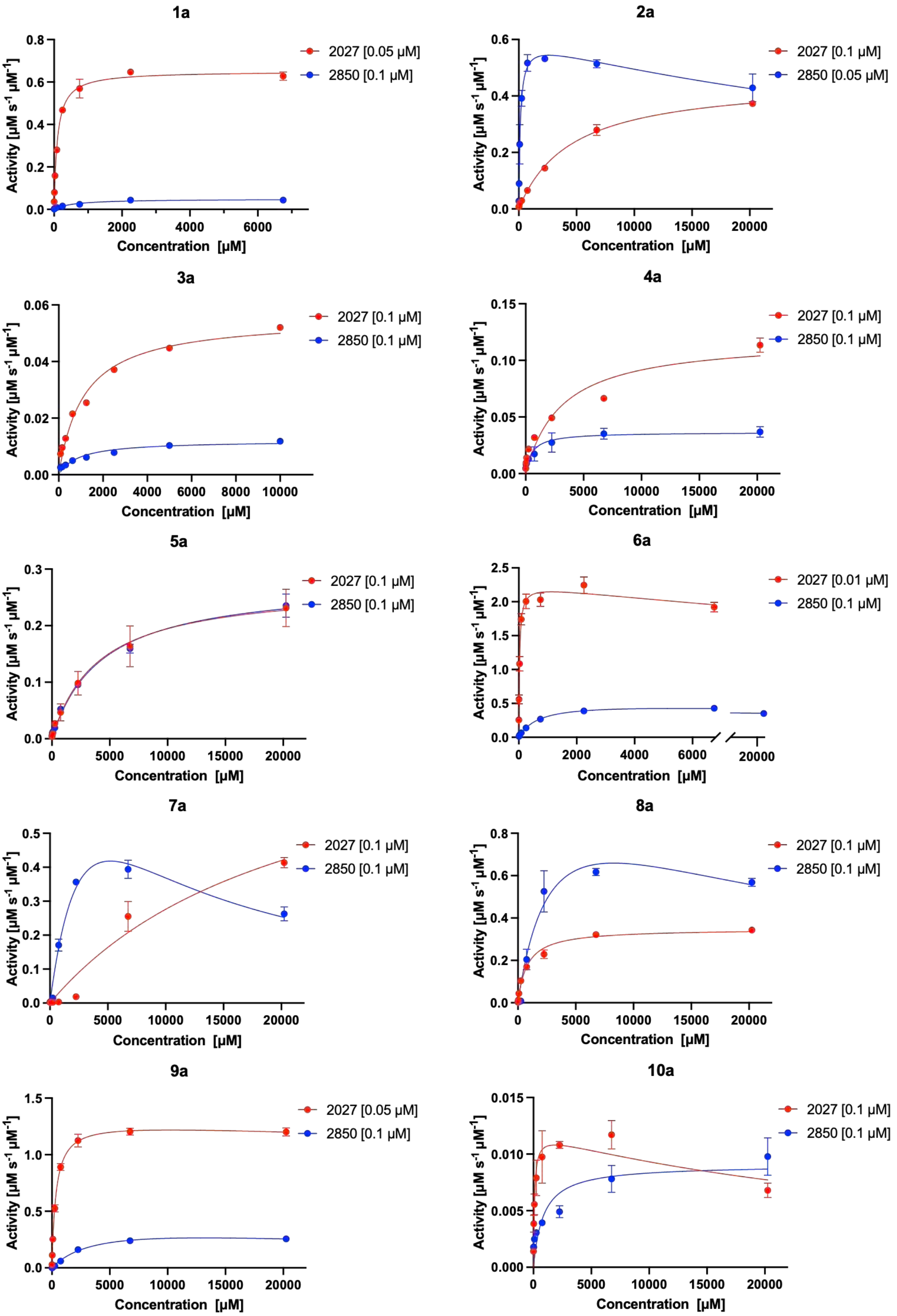


**Figure S3:** Michaelis-Menten kinetics graphs. Kinetic assay was performed using 50 mM tris buffer (pH 8.0), 20 mM $F_{420}$, 20,250-3.08 mM substrates, 0.01-0.1 mM enzyme.

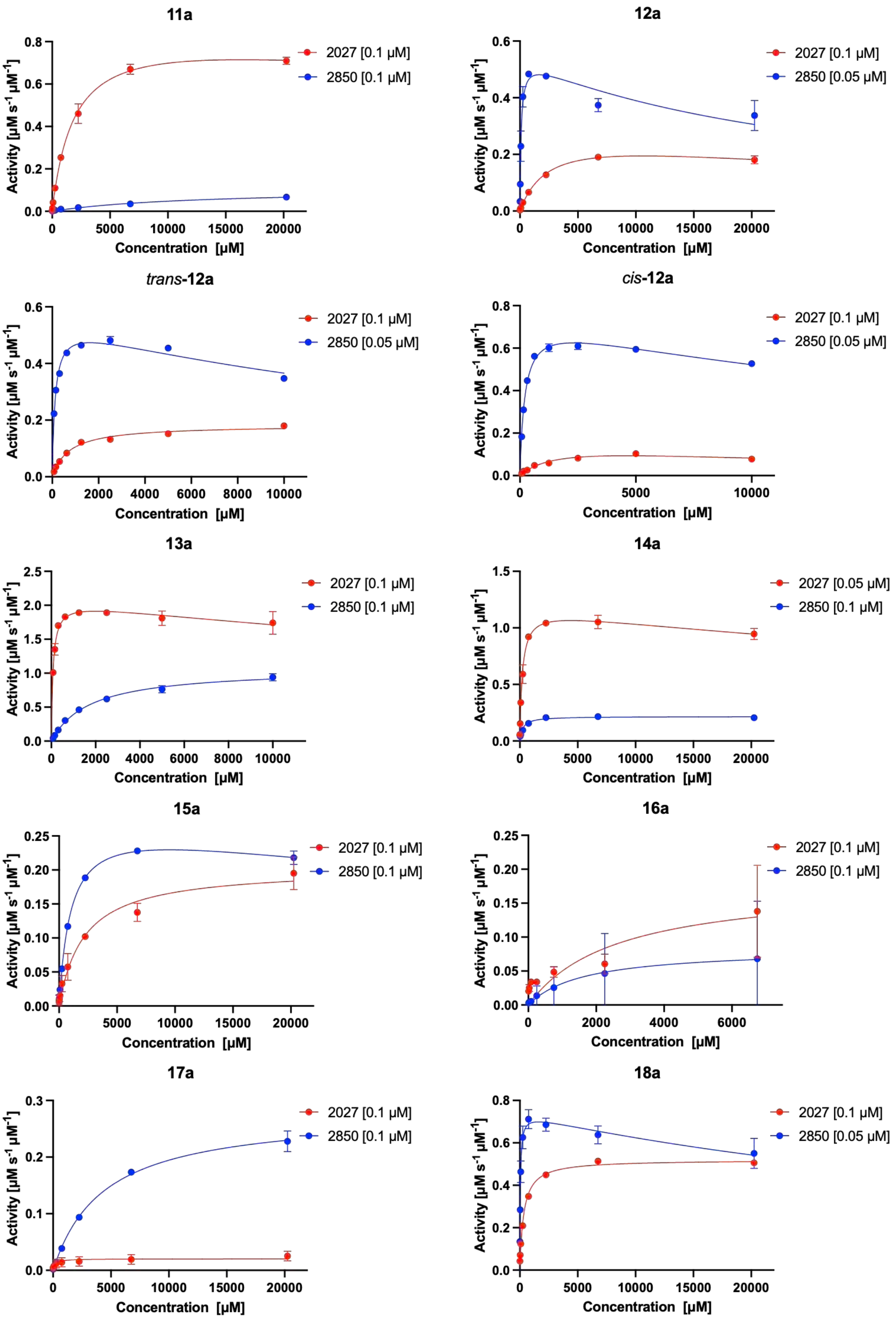


**Figure S3 (cont.):** Michaelis-Menten kinetics graphs. Kinetic assay was performed using 50 mM tris buffer (pH 8.0), 20 mM $F_{420}$, 20,250-3.08 mM substrates, 0.01-0.1 mM enzyme.

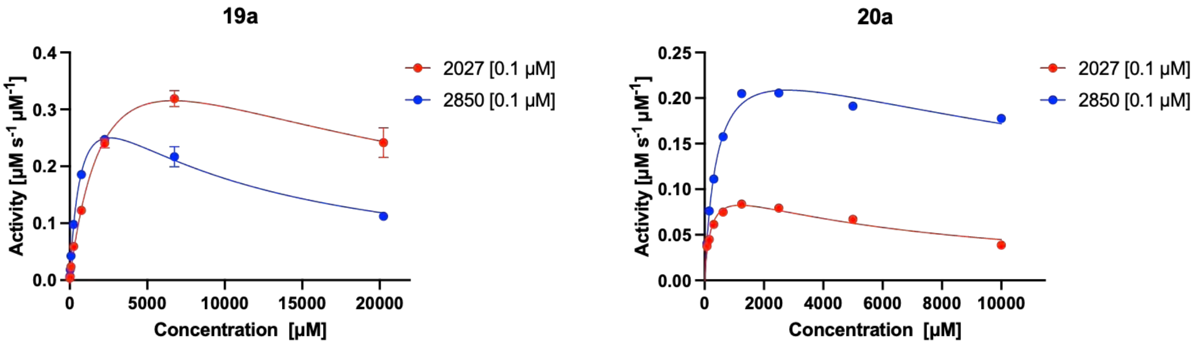


**Figure S3 (cont.):** Michaelis-Menten kinetics graphs. Kinetic assay was performed using 50 mM tris buffer (pH 8.0), 20 mM $F_{420}$, 20,250-3.08 mM substrates, 0.01-0.1 mM enzyme.

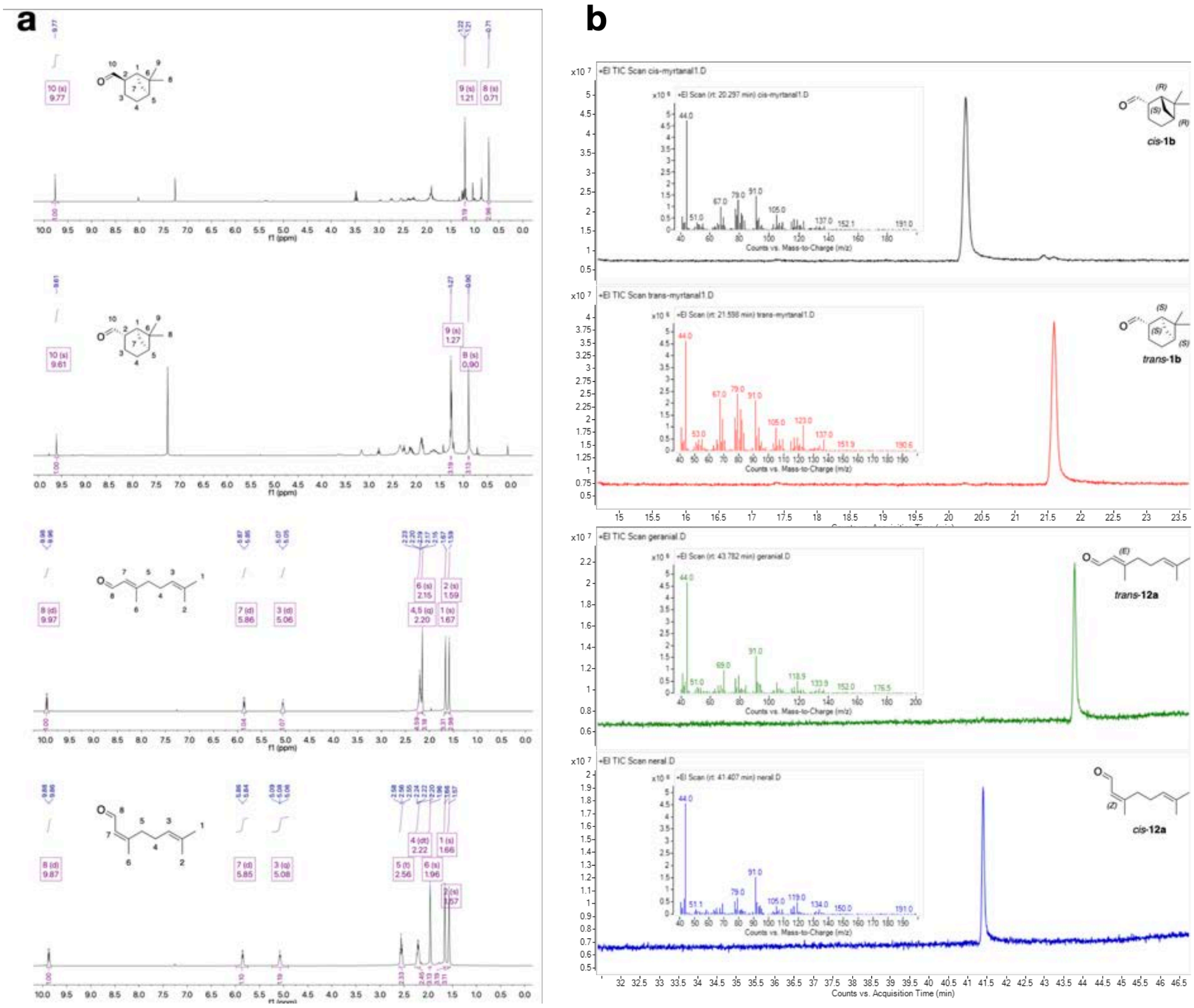

**Figure S4.** Confirmation of products from Dess-Martin oxidation. **a** [1]H NMR spectrum. **b** GC-MS chromatogram and MS spectrum (inset) of *cis*-, *trans*-**1b** and *cis*-, *trans*-**12a**.

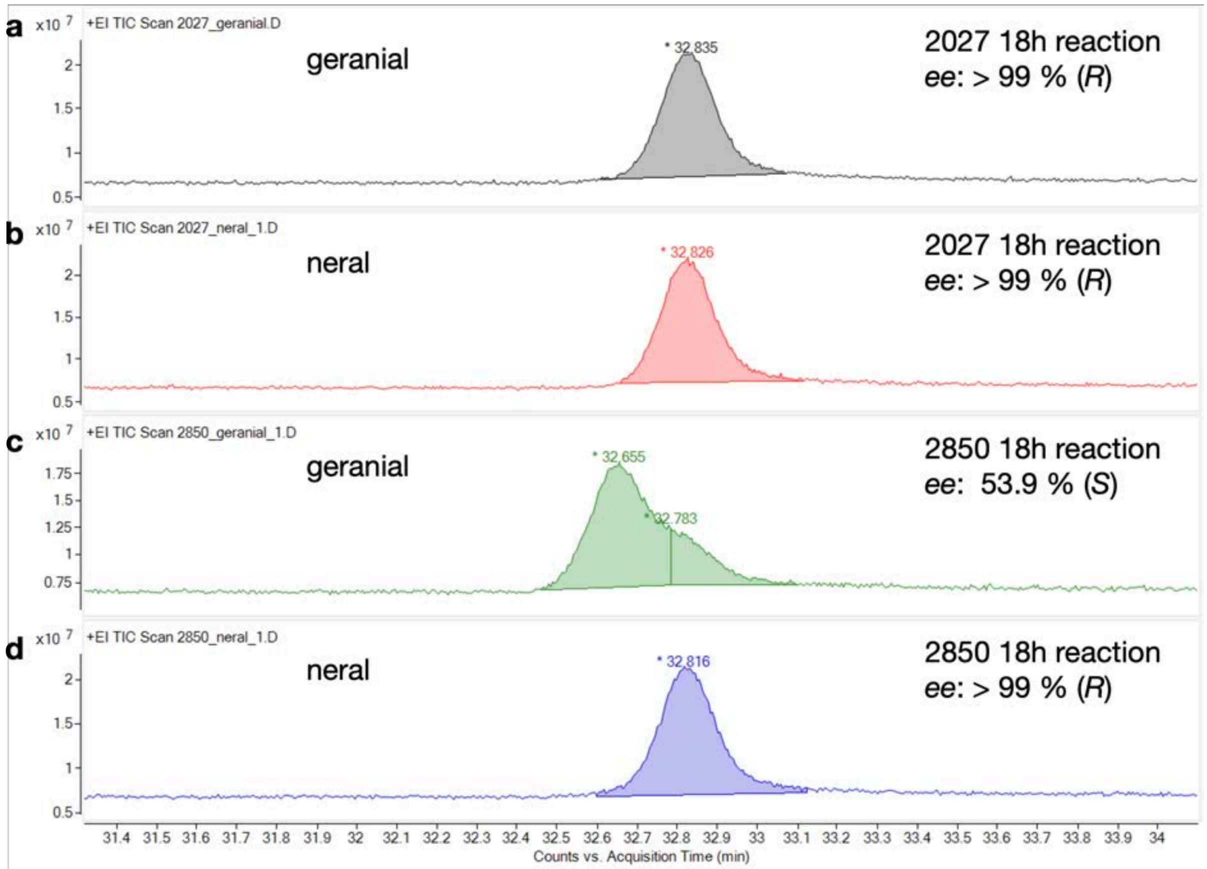


**Figure S5.** Chiral GC of reduction of geranial (**a**, **c**) and neral (**b**, **d**).

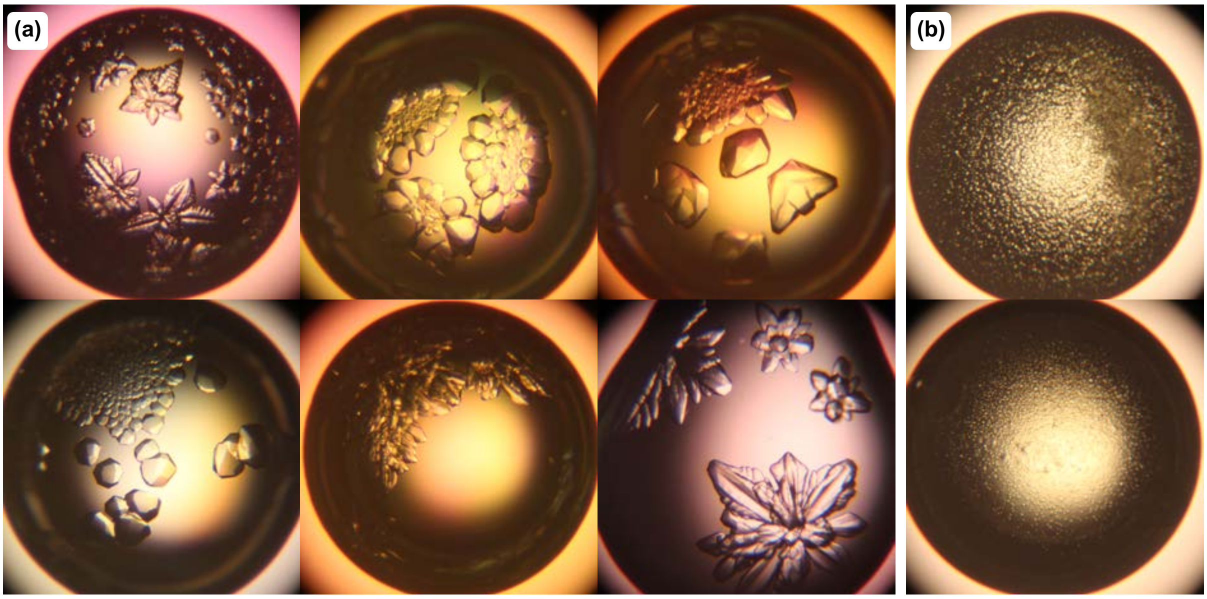


**Figure S6.** Crystallization of MSMEG_2850. **a**. Apoenzyme crystals (Sodium phosphate monobasic monohydrate:0.67~0.77M, potassium phosphate dibasic 1.25~1.4 M, pH 6.9), **b**. Attempts to form microcrystals of $F_{420}$-bound complex

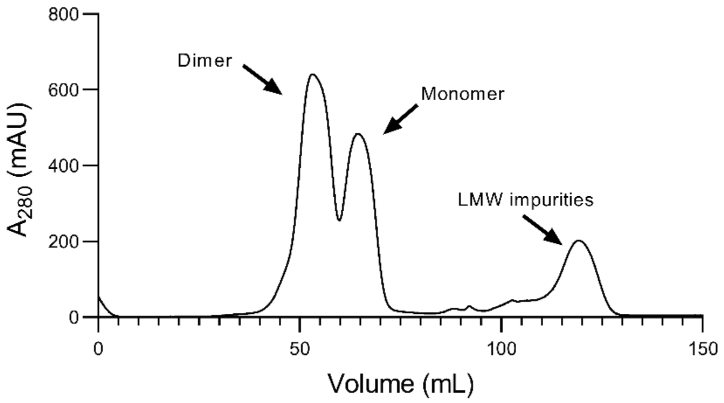


**Figure S7:** Gel filtration of MSMEG_2850 in the absence of $F_{420}$. Sample was run on a HiLoad 16/60 Superdex 75 pg (GE Healthcare) equilibrated with 20 mM HEPES, 150 mM NaCl, pH 7.5. Two distinct peaks with $R_v$ of 53 mL and 65 mL corresponding to the dimer and monomer, respectively, are observed.

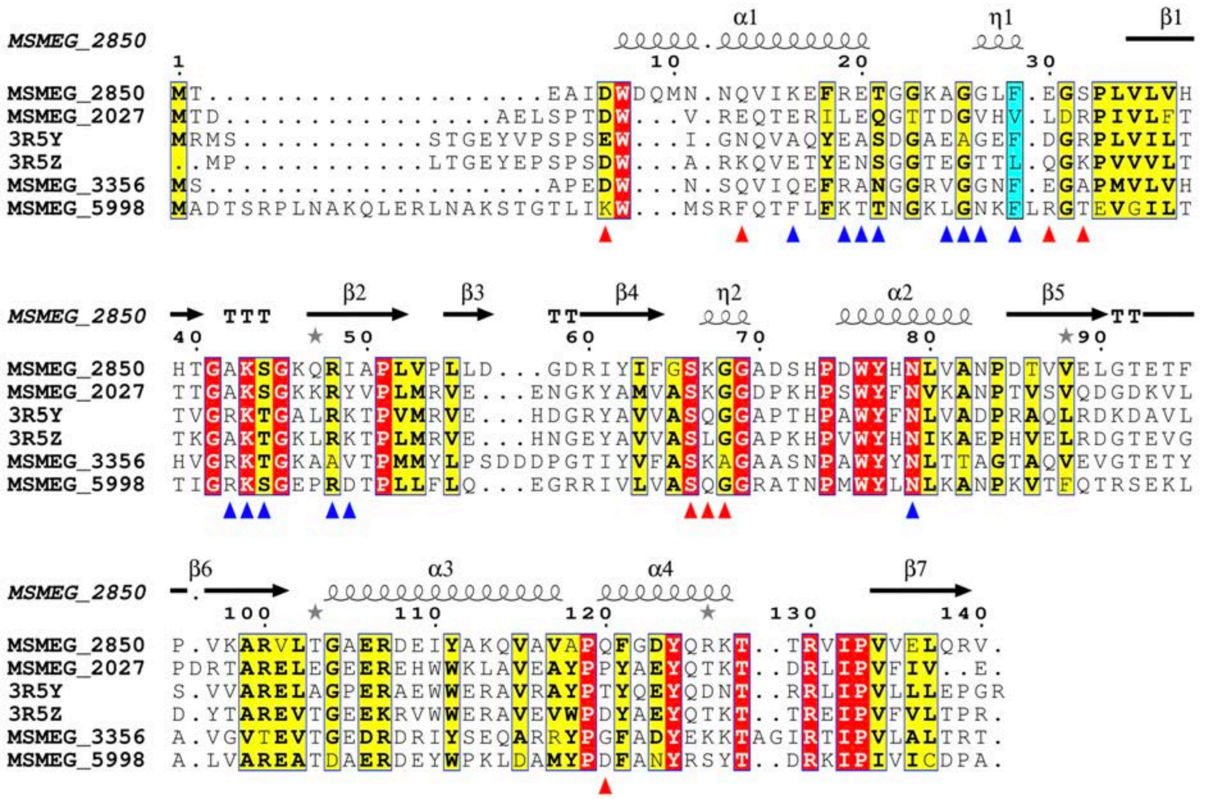

**Figure S8:** Multiple sequence alignment of FDOR-A1s. Sequences were aligned as per materials and methods. Positions involved in hydrogen bonding or salt bridges in the dimer interface of MSMEG_2027 and MSMEG_2850 are indicated by red and blue triangles, respectively. Despite many interface residues being highly conserved between all the sequences the sets of interface residues involved in polar contacts do not intersect between the two structures.

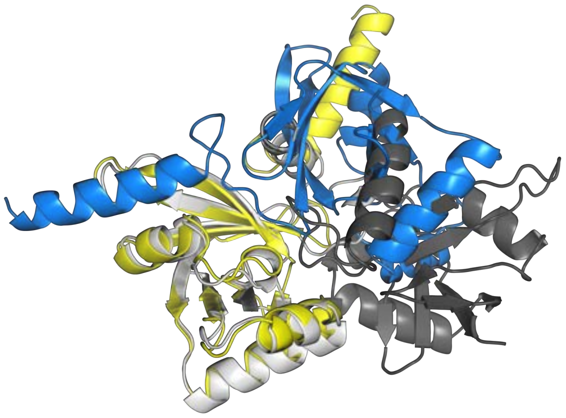

**Figure S9:** Structural alignment of the biological dimers of MSMEG_2027 (6XRI) and MSMEG_2850 (8D4W). The relative orientation of each monomer differs between the dimers of MSMEG_2027 (yellow and blue) and MSMEG_2850 (black and white) and thus the dimers are not superimposable. This is consistent with the lack of any symmetry operations common to both the $I4_132$ and $P2_1$ space groups (other than the trivial identity operation).

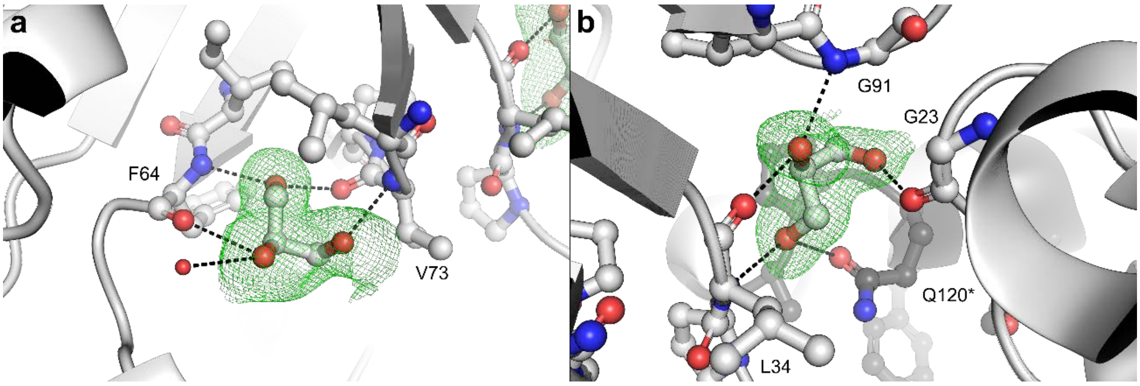

**Figure S10:** Polar interactions of cocrystallized glycerol molecules with MSMEG_2850. Polder omit maps contoured to 3σ around the ligands are shown. a Interactions between Gol201 b Gol202 are shown as dashed lines.

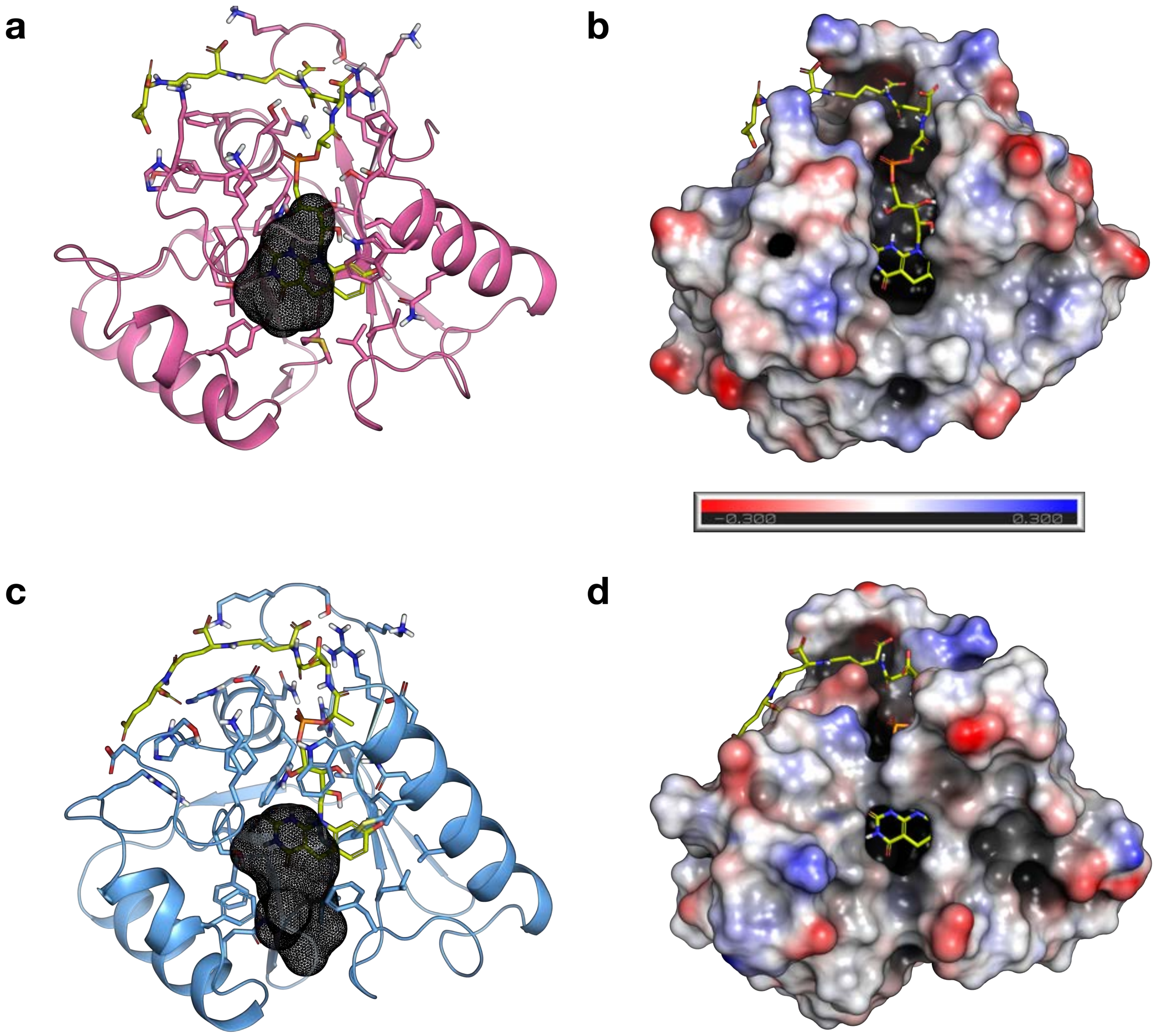


**Figure S11** Complex models with $F_{420}$ bound. MSMEG_2027 is modelled from a holoenzyme structure (PDB 6WTA) (**a**, **b**). The apoenzyme structure solved in the study is used to model MSMEG_2850 (PDB 8D4W) (**c**, **d**). Residues within 5 Å from $F_{420}$ are displayed as sticks, calculated binding site volume are shown in black meshes (**a**, **c**). Electrostatic potential maps are shown between -0.3 (red) and +0.3 (blue) in panels **b** and **d**.

a

b

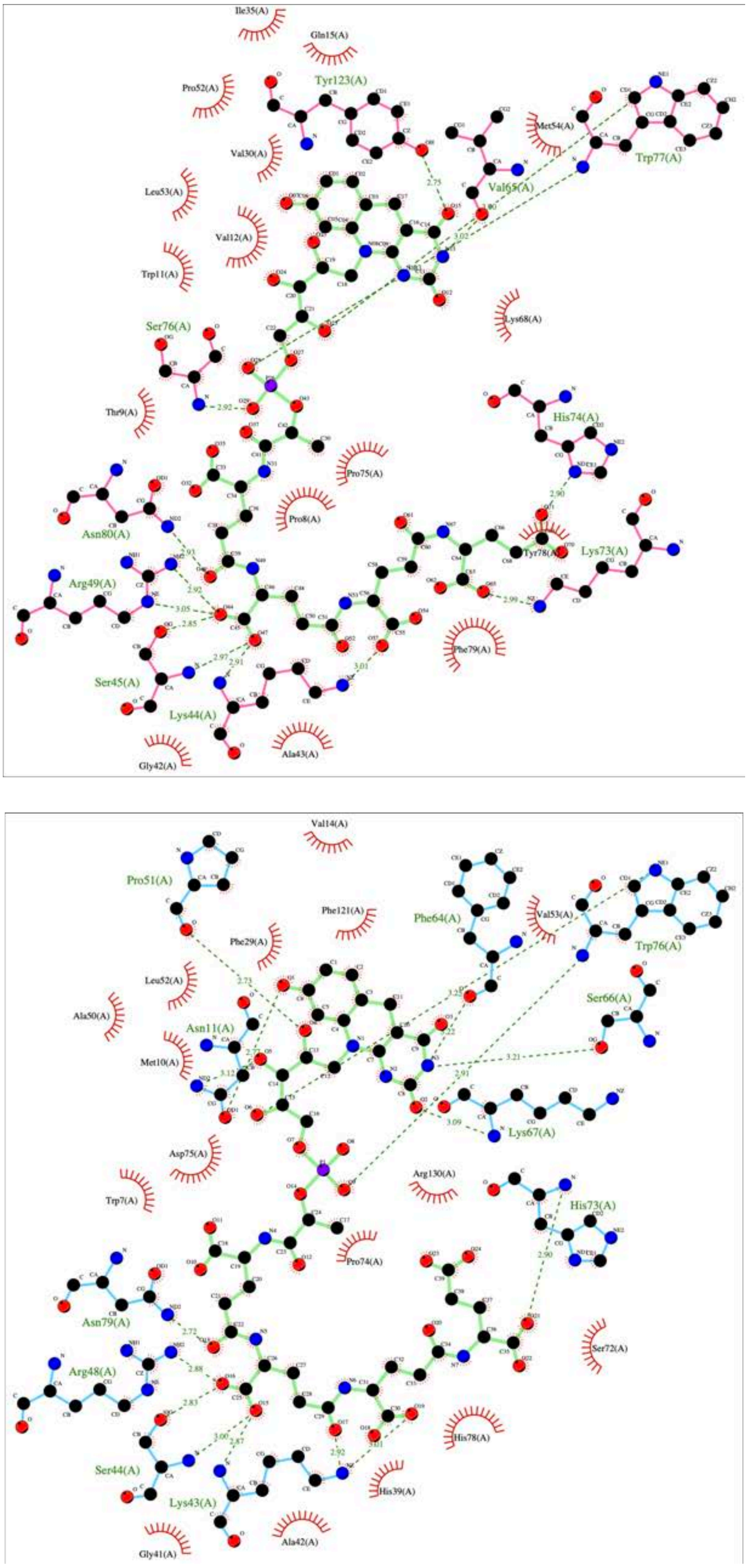


**Figure S12**: Interactions between $F_{420}$ and the active sites of MSMEG_2027 (**a**) and MSMEG_2850 (**b**) were examined using LigPlot plus.[6] Hydrophobic contacts and hydrogen bond interactions are represented by "eyelashes" and dashed lines, respectively.

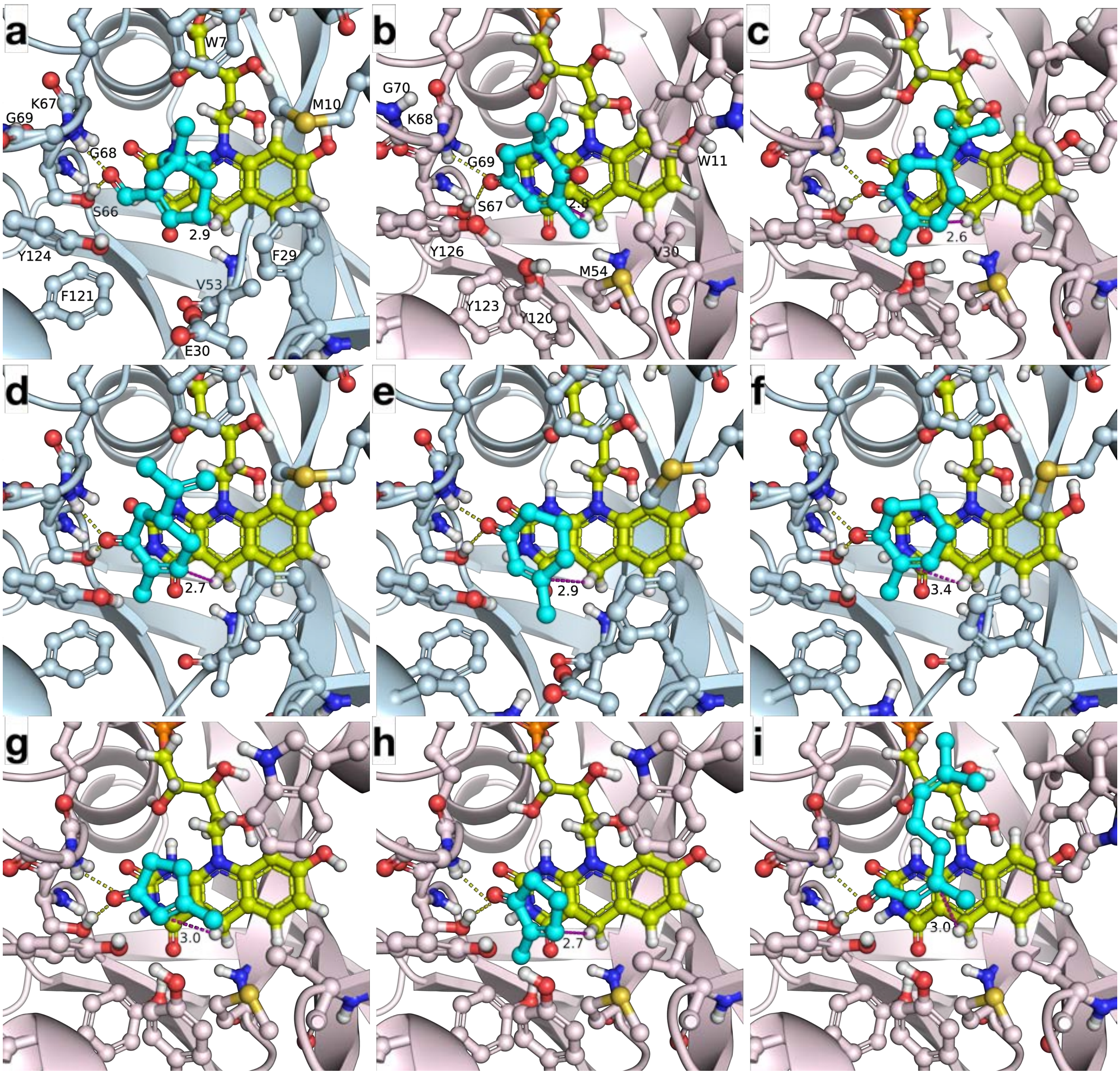


**Figure S13.** Binding modes of prochiral substrates that are not shown in Figure 6. MSMEG_2027 and MSMEG_2850 are coloured pink and blue, respectively. Panel a, **1a**. Panel b, **2a**. Panel c, **4a**. Panel d, **5a**. Panel e, **7a**. Panel f, **8a**. Panel g, **10a**. Panel h, **11a**. Panel i, *trans*-**12a**. Hydrogen bonds are shown as yellow dashed lines. The distance between the hydrogen of C5 in $F_{420}$ and carbon atom of substrate aligned for hydride transfer is shown in Å with a magenta dashed line. Substrates and $F_{420}$ appear in cyan and yellow green, respectively.

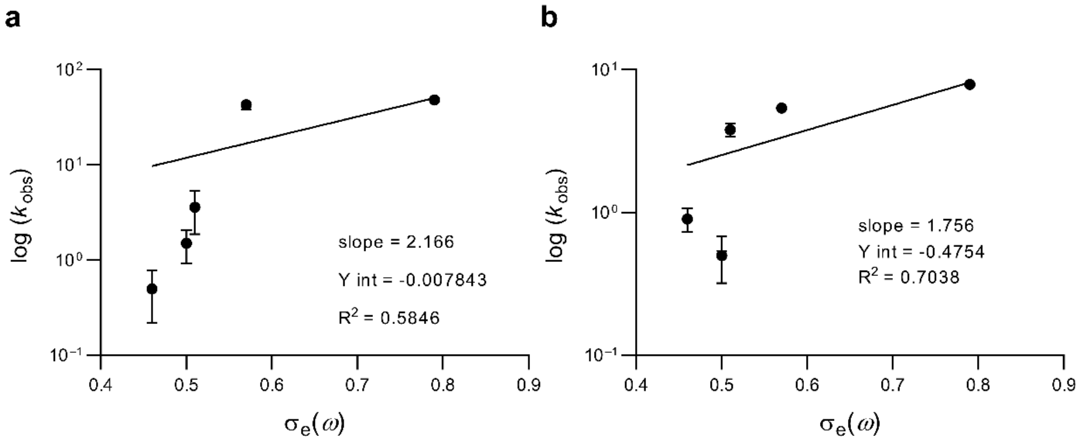


**Figure S14.** Effect of intrinsic electronic component $\sigma_e(\omega)$ of the Hammett $\sigma_p$ of the electron-withdrawing group of substrates **13a–17a**. A general positive correlation is observed between EWG strength and activity. Activities of MSMEG_2027 (**a**) and MSMEG_2850 (**b**) are shown. Data from **Table S4**.
